# Modelling temporal shift-invariance in self-supervised generative models improves accuracy and interpretability of species detection in soundscape recordings

**DOI:** 10.64898/2025.12.09.693207

**Authors:** K. A. Gibb, A. Eldridge, A. L. Shuaibu, I. J. A. Simpson

## Abstract

Realising the potential for acoustic monitoring to deliver biodiversity insight at scale requires new approaches to the automated analysis of PAM recordings that are trustworthy as well as cost-effective. Discriminative models trained on annotated species data are gaining popularity but are labour intensive, notoriously opaque and biased. Self-supervised generative models such as Variational Autoencoders (VAE) offer great potential for learning compact yet expressive representations of data, which can be used for subsequent discriminative tasks and are intrinsically interpretable. However, the default learning algorithm results in weakly discriminative data representations due to under-specification of the generative task. We propose and evaluate a novel modification to the VAE learning algorithm that models intra-frame shift-invariance. We demonstrate that this modification provides representations that are more interpretable, consistent and improve classification performance. Performance accuracy is evaluated on species detection tasks on two weakly annotated data sets across temperate and tropical terrestrial habitats and compared to leading discriminative models BirdNet and Perch, as well as the classic VAE. Whilst demonstrated in terrestrial recordings, the approach is transferable to marine, freshwater, and soil habitats. These innovations set the path for trustworthy, data and time-efficient tools to support solid ecological inference from large-scale passive acoustic monitoring surveys.

## 1 Introduction

In the face of escalating biodiversity loss, Passive Acoustic Monitoring (PAM) has been heralded as a potentially transformative biodiversity monitoring tool [1]. To meet this potential, the ease with which data can be collected needs to be met with efficient, interpretable and accurate models that support solid ecological inference. Two basic approaches dominate in Passive Acoustic Monitoring: species identification, and soundscape description. The former is fast growing in popularity thanks to discriminative models such as BirdNET [2], but despite rapid uptake, issues of reliability undermine trust; the latter has come under criticism due to lack of reliable, interpretable, generalisable means to describe whole soundscape recordings [3]. A lack of theories in ecoacoustics means it is critical not to fall intowpurely data driven approaches; in application, environmental changes are long term, so confidence is needed in evidence-based decision making processes. Here we propose a third way that integrates the efficiency and ability to encode prior ecological knowledge of first generation soundscape indices, with the scalable, and generalisable datadriven characteristics of deep learning, whilst delivering a route to interpretability.

The boom in machine learning for PAM has been driven in part by the ready availability of large crowd-sourced labelled audio data such as [eBird](https://ebird.org/), [iNaturalist](https://www.inaturalist.org/) and Xeno Canto enabling supervised training of deep learning models for avian species. Growing repositories for invertebrate, anuran, marine and freshwater species suggest respective models may soon follow. BirdNet and Perch [2, 4] demonstrate reasonable performance in both specific [5] and more general tasks [4], but suffer fundamental challenges. These include: bias due to non-representative data sets that limit generalisation [6, 7]; unreliable performance in ‘noisy’ settings and where acoustic diversity is high (i.e. in dense choruses or otherwise crowded soundscapes that are the norm rather than exception) [8, 9]; poorly calibrated detection thresholds [10] and overly confident misidentification of out-of-distribution (OOD) signals [11]. These combine to create real world issues such as presence of water sounds leading to confident identification of water bird species when there are none.

Additional data labelling and innovations in data augmentation and learning strategies go some way to addressing problems such as under-represented classes [12], but unless a sufficient number and diversity of precisely labelled (i.e. timestamped) data are available to identify the learning task [13], these issues will persist. In terms of interpretability, methods for visual interrogation of model predictions such as class activation maps (CAMs) exist [14] but their application has yet to be convincingly demonstrated in whole soundscape recordings and are not currently supported by BirdNET. Heinrich et al [15] examine a method for identifying and interrogating bird call prototypes, providing a novel interpretability mechanism. However this approach relies on a fixed number of prototypes for known classes, potentially neglecting or confusing important outliers. Prototypes themselves cannot be directly examined — a limitation we address — and it is yet to be tested in dense soundscapes. Architectures that are inherently interpretable, as well as cost-effective to train, are critical to build trust in PAM as a transformative ecological macroscope.

Self-supervised methods offer an attractive alternative to create cost-effective, generalisable and interpretable models. Self-supervised learning results in *general* representations from large-scale data. By learning a representation of the whole data distribution of the soundscape we preserve more of the relevant source signal in contrast to using embeddings from a discriminative model trained to identify specific instances. This should act as a *strong and relevant prior* to aid inference of species presence, particularly for weakly labelled data. This reduces the need for strong human annotations and enables more cost-effective approaches. Furthermore, by learning the whole distribution, self-supervised methods tend to result in models that are more flexible [16]. We propose they are therefore better suited to modelling complex soundscapes (where multiple species vocalisations mix with geophony and anthrophony etc.) than the discriminative models used for species identification to date, whilst still affording potential for species classification. Thirdly, self-supervised architectures afford the opportunity to build inherently *interpretable* models using paradigms such as disentangled representation learning [17, 18]. Despite growing success in image audio and speech processing (see [19] for a review), un- and self-supervised deep learning approaches have yet to be fully developed for soundscape analysis (for some excellent early work in this area see [20, 21, 22, 23, 24, 25, 12]). In this article we propose that self-supervised learning in general, and Variational Autoencoders (VAEs) [26, 27] in particular, offer an underexplored opportunity to develop cost-effective, flexible and interpretable soundscape representations that have potential to address several key challenges of current discriminative approaches. We suggest here that VAEs offer a valuable alternative to the discriminative models that are used for species ID and heuristic soundscape descriptors, one that provide an efficient means to learn representations for classification, clustering or detection tasks.

In [28] we introduced a visualisation mechanism for VAEs that demonstrated interpretability for ecoacoustic classification for the first time. Whilst VAEs are intrinsically interpretable and data efficient, there are drawbacks in the classic implementation that reduces their performance of learned representations in downstream classification or detection tasks. Due to the ill-posed nature of self-supervised representation learning in complex multi-modal data such as raw soundscape audio, representations tend to become entangled, mixing together the underlying factors of variation [29]. For example, when training a VAE the spectrogram is spliced into consecutive frames and the algorithm learns a compact representation of each frame in a latent space. Note that the *absolute* temporal position of events that lie within that frame (e.g. the onset of a particular bird vocalisation) is arbitrary when frames are segmented automatically (as is the norm). However, during training the classic VAE algorithm finds the most parsimonious compressed latent representation that minimises the difference between input and generated spectrograms. Spurious information (such as distance from frame start to signal onset) carries as much weight as more ecologically meaningful information (such as the distance between peaks or nuances of spectral morphology which represent acoustic traits of a given species). To reduce this effect, prior knowledge and heuristics can be introduced as inductive bias and regularisation in order to encourage latent representations to describe the key signal characteristics of interest with greater consistency [30]. Such approaches have yet to be explored for modelling natural soundscapes.

Here we propose a method to increase relevance of learned representation by embedding task-specific prior knowledge to structure the latent space. On the basis that *when* a vocalisation occurs within a given frame is essentially arbitrary, we introduce a representation scheme that is invariant to the absolute onset time of sound events within a given frame. We call this intra-frame *shift invariance*. Using terrestrial soundscapes as a test case, we evaluate the impact of these interventions on both discriminative accuracy and model interpretability in a weakly labelled species detection task in comparison to gold standard discriminative models. Our results show that the shift-invariant modification yields improved discriminative performance in a species detection task; shift-invariant VAEs out perform the classic VAE architecture and are competitive with large discriminative models, whilst requiring a order of magnitude less training data. Importantly this method provides a novel interpretability pipeline for interrogating the basis of a species detection.

## 2 Results and Discussion

In this section our adjustments to the learning algorithm are evaluated in terms of accuracy at species detection and improvement in the interpretability of generated representations. We set two different models: (1) a classic VAE and (2) an intra-frame shift-invariant VAE (SIVAE) alongside the current openly available gold standard discriminative models BirdNET and Perch [2, 4]. We begin by outlining the datasets before cross-examining the performance and interpretability provided across model classes.

### 2.1 Datasets

#### 2.1.1 Sounding Out (SO)

The Sounding Out dataset, originally described in [31] comprises large-scale acoustic surveys carried out across a gradient of ecological degradation in Ecuador and the United Kingdom. Each audio sample is 60s and recorded at a sample rate of 48kHz. The dataset is approximately balanced across countries, with 1620 and 2160 samples for training and 405 and 540 samples for testing (80/20) from the UK and Ecuador, respectively, with both containing regular occlusions as samples are drawn from the dawn and dusk choruses.

Avian species richness and abundance estimates for each recording were collapsed to binary per-species presence/absence indicators. Estimates provided by local experts of the density of biophony within the soundscape and annotations indicating the presence and degree of wind and rain were used to manually select a test subset that exhibit common occlusions, i.e. high overall biophonic density, avian abundance and/or dense wind or rain. We describe this in figures as the ‘occlusions’ subset; it consists of 16 samples from each site. The full test set contains this subset plus randomly sampled observations up to a proportion of 0.2. The label frequency distribution for each country is long-tailed with very few observations for certain species. We elected to train only on species with greater than 10 presence labels to ensure enough signal for learning while preserving most rarer species. This leaves at total of 46 bird species in the UK where 40% have fewer than 50 training examples, and 90 bird species in Ecuador where 66% have fewer than 50 training examples. For a full description including survey and annotation methods see [31]. Species labels counts are provided in Figure 10.

#### 2.1.2 Rainforest Connection (RFCX)

The RFCX dataset, originally described in [32], is the training set from the 2021 Kaggle Species Audio Detection challenge. Recent research fine-tuned BirdNet and Perch on this dataset using a few-shot learning approach [4]; this provides a comparator for a species presence prediction task. RFCX contains Puerto Rican soundscape recordings recorded at 60s at a sample rate of 48kHz with annotations for 21 avian and anuran species. RFCX is similar in size to SO, with 3782 samples for training and 945 samples for testing (80/20). Species specific song labels were extracted using a template matching algorithm providing spectro-temporal bounding boxes around calls that were manually verified in [32]. Under the assumption that lack of a bounding box implies absence, we collapse strong labels to a binary presence / absence flag for each species. This leaves 13 bird species where 90% have fewer than 50 training examples and 11 frog species where all have fewer than 50 training examples. The audio contains some occlusions of species calls, though not to the same degree as SO. For a full description of the dataset including survey and annotation methods see [32].

#### 2.1.3 Dataset limitations

Both datasets feature a certain amount of label noise, resulting in a more challenging task. In the case of the SO UK dataset, we found through observation that certain species were occasionally misidentified. For the RFCX which were extracted using a template-matching approach there are many examples that contain unlabelled vocalisations for different species due to natural variability in calls. We note however that this scenario is realistic in the context of PAM and and aligns with our aim of providing a more cost-effective pipeline using weak labels.

### 2.2 Model setup and evaluation methods

VAEs were pre-trained to embed and reconstruct log mel spectrograms before downstream linear classifiers were fit to samples from the learned posterior to predict weak presence / absence labels for each dataset independently. Performance is compared with (i) out-of-the-box BirdNET V2.4 and (ii) the latest Perch and BirdNet few-shot learning results published by Ghani et al. [4]. The full details of the CNN architecture of VAEs can be found in our previous work [28] while refinements and training paradigms relevant for this study are described in 4.

Detailed descriptions of score metrics are detailed in 4.4.2 but to briefly summarise, detection performance is evaluated use three common score metrics: (i) the average precision (AP), (ii) the area under the receiver operating characteristic curve (auROC) and (iii) Top-1 scores. Aggregate results are shown in Table 1 and full score distributions in 2.

**Table 1:** Area under the ROC (auROC), mean average precision (mAP) and top-1 accuracy scores for species detection models trained on latent space representations for each model variant alongside BirdNET and Perch. Scores are averaged across the entire species community over 3 different training runs for each dataset. Comparisons are made with both out-of-the-box BirdNET V2.4 and the best results from Ghani et al. [4] where BirdNet and Perch were fine-tuned on a few-shot learning task. * indicates “strong labels” where bounding boxes were used to delineate the call in the training data. Size denotes the dimensionality of the embedding for the corresponding temporal resolution in seconds.

| Model | Size | Duration (s) | SO Ecuador |  |  | SO United Kingdom |  |  |
| --- | --- | --- | --- | --- | --- | --- | --- | --- |
|  |  |  | auROC | mAP | Top-1 | auROC | mAP | Top-1 |
| VAE | 128 | 1.536 | 0.86 (0.002) | 0.36 (0.001) | 0.50 (0.012) | 0.77 (0.004) | 0.33 (0.004) | 0.58 (0.006) |
| SIVAE | 128 | 1.536 | 0.87 (0.002) | 0.32 (0.003) | 0.47 (0.002) | 0.81 (0.002) | 0.38 (0.006) | 0.61 (0.006) |
| BirdNET V2.4 | 1024 | 3 | 0.86 | 0.42 | 0.41 | 0.82 | 0.46 | 0.50 |
|  |  |  | RFCX Birds |  |  | RFCX Frogs |  |  |
|  |  |  | auROC | mAP | Top-1 | auROC | mAP | Top-1 |
| VAE | 128 | 1.536 | 0.88 (0.014) | 0.25 (0.026) | 0.49 (0.084) | 0.89 (0.003) | 0.22 (0.013) | 0.52 (0.021) |
| SIVAE | 128 | 1.536 | 0.92 (0.002) | 0.34 (0.006) | 0.67 (0.045) | 0.90 (0.001) | 0.24 (0.004) | 0.54 (0.016) |
| BirdNET V2.4 | 1024 | 3 | 0.89 | 0.26 | 0.52 | - | - | - |
| BirdNet V2.2 (few-shot*) [4] | 1024 | 3 | 0.96 | - | 0.79 | 0.96 | - | 0.75 |
| Perch (few-shot*) [4] | 1280 | - | 0.97 | - | 0.83 | 0.96 | - | 0.75 |

Model interpretability is assessed by making visual comparisons between real prototypical calls of target species and spectrograms generated by shifting across the decision boundary for each species detection model as described in [28]. This provides direct inspection of the spectro-temporal features of species-specific calls identified in feature space. Species-specific scores are shown alongside generated calls to highlight the difference in discriminative ability alongside model interpretability. We identify, discuss and suggest methods to combat significant failure cases for future work.

### 2.3 Evaluating performance against large discriminative models in a species detection task

In this section we examine the differences in performance between VAE representations and BirdNET and Perch on a multi-label species presence detection task. Table 1 and Fig. 2 show a comparative breakdown between the classic VAE and the SIVAE, along with BirdNet out-of-the-box, and fine-tuned BirdNet and Perch as described in [4]. Table 1 details mean average precision, area under the ROC curve and top-1 scores while 2 shows the full distributions of each metric.

**Figure 1.**
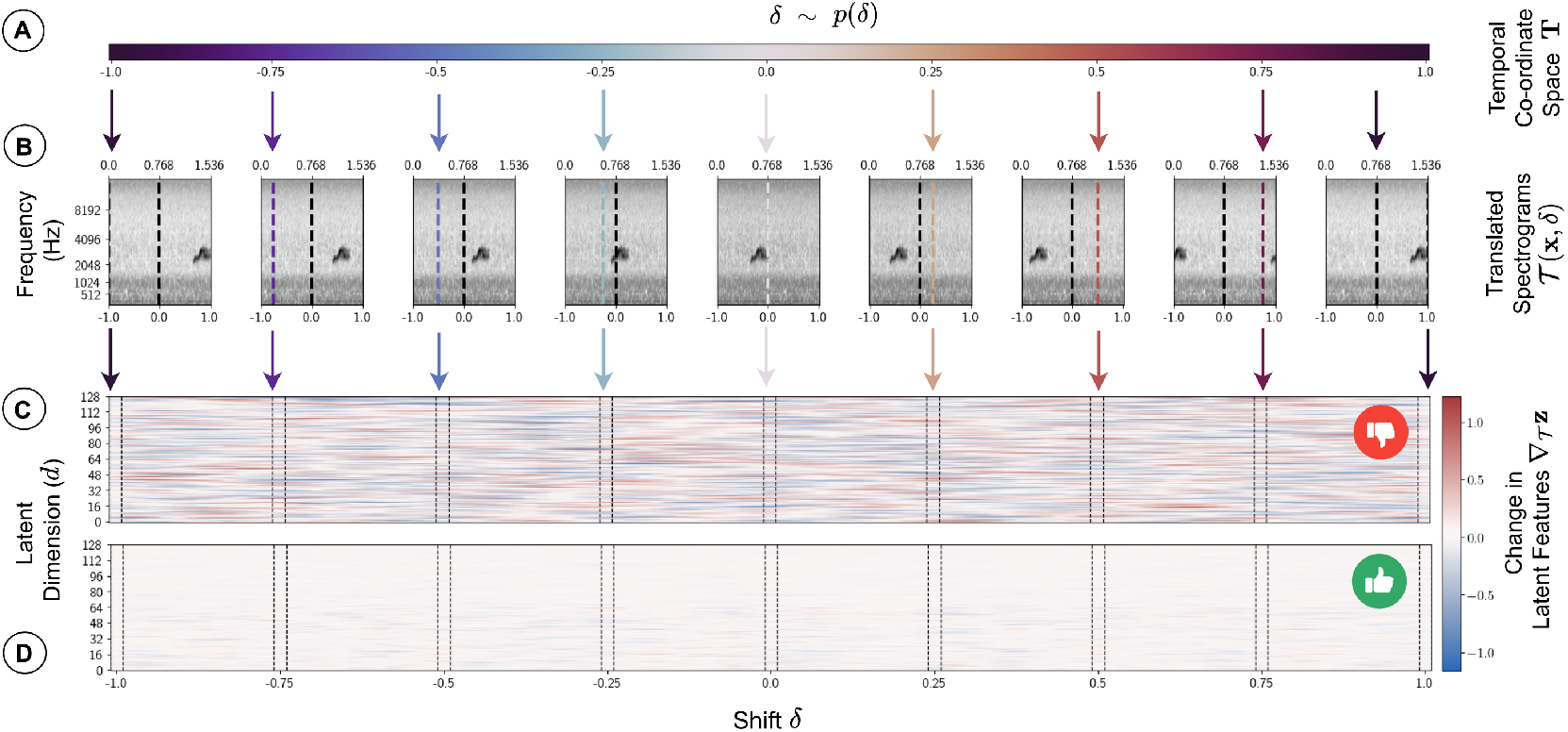
A demonstration of how the VAE learns representations **z** that are less consistent than a shift-invariant model. **(A)** A classic VAE implicitly encodes an arbitrary reference co-ordinate system *T* denoting the absolute position of the signal in time within a framed spectrogram **x. (B)** Applying a shift *δ* via a translation operator 𝒯 (**x**, *δ*) forward or backwards in time fundamentally describes the same information, the spectro-temporal morphology of the bird call, regardless of where it lies with respect to the co-ordinates. **(C)** A classic VAE representation moves around in the latent space as the temporal shift is applied: it is not a consistent description of the bird call itself. **(D)** By disentangling infra-frame shift, latent space representations are invariant to the absolute position of the signal resulting in a more consistent description of the spectro-morphology of the call.

**Figure 2.**
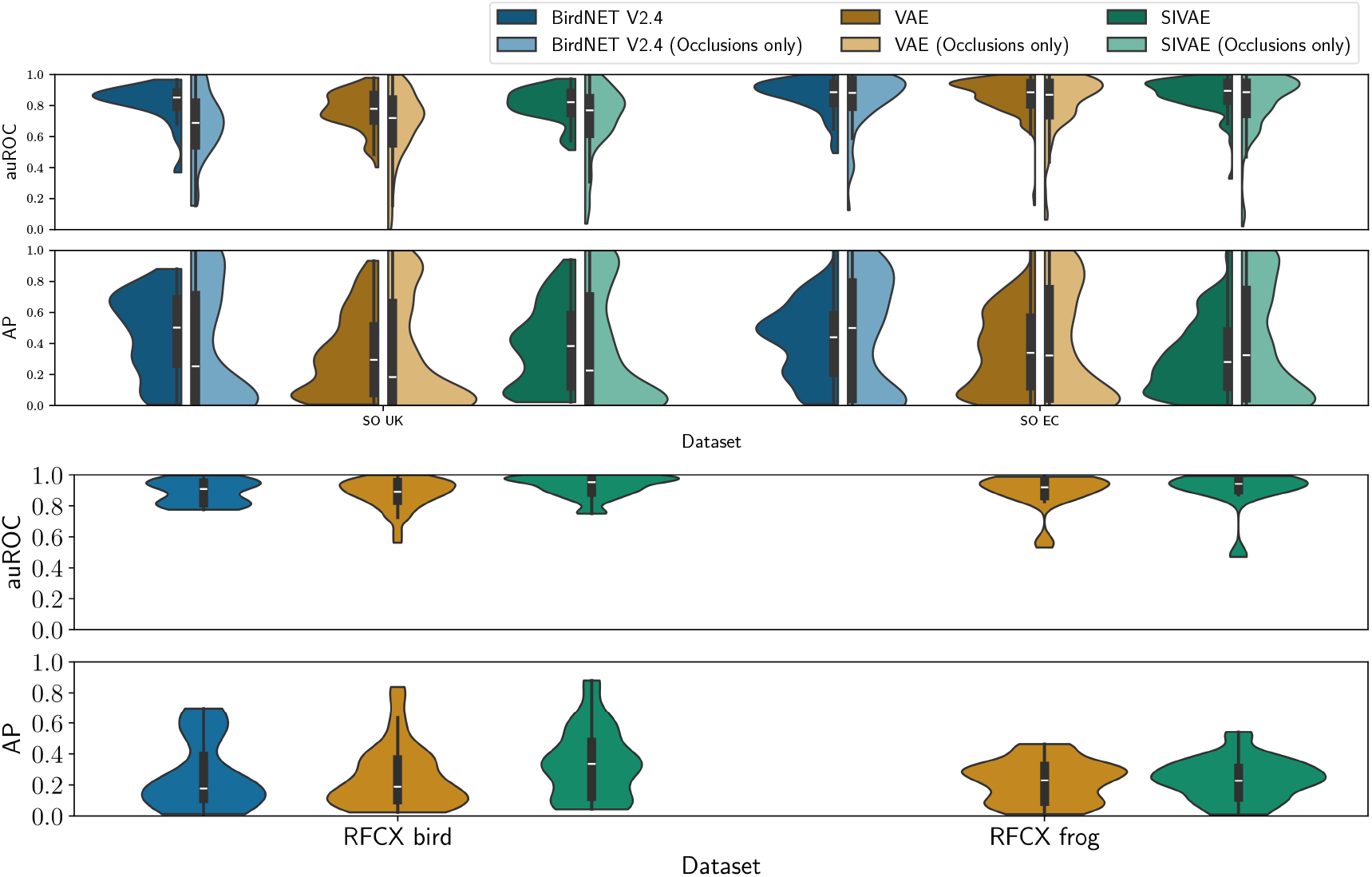
Area under the ROC curve (auROC) and average precision (AP) score distributions across out-of-the-box BirdNet V2.4 and both VAE variants. Full distributions show across species the auROC is approximately equivalent with BirdNET and across the VAE and SIVAE for the full dataset, while our models typically perform slightly better on examples with high occlusions. Both VAE variant representations provide less precision than BirdNET in the SO dataset. Introducing shift-invariance results in a small improvement in the UK and a drastic improvement in RFCX bird dataset, while we see a slight drop in precision in Ecuadorian soundscapes.

Overall, performance is comparable with BirdNET V2.4 despite the use of very weak labels, far fewer training examples and a higher compression factor. SIVAE embeddings demonstrate high quality species detection, with maximum scores on RFCX (0.92 auROC, 0.34 mAP) achieving higher than BirdNET (0.89 auROC, 0.26 mAP). Significance testing using the Wilcoxon test shows a drop in mAP in SO (UK −0.08015, p-value 0.012163, EC −0.09969, p-value 0.00016) with no significant difference in auROC. For RFCX, we achieve a slight improvement auROC (0.043341 p-value 0.03222) with no significant difference in mAP (0.09061, p-value 0.10156). Given the difficult nature of the task, we achieve respectable performance on SO EC and SO UK, suggesting this approach has a high quality discriminative ability for the presence of most species individually even in the presence of dense chorusing where segmenting distinct calls from the cachophony is no easy task. Figure. 2 shows VAE and SIVAE representations fail to achieve as high AP as BirdNET in the SO dataset though a clear improvement is made in the UK between VAE models by introducing shift-invariance. In this case, at the upper end of the distribution, SIVAE representations are slightly less discriminative than BirdNET, while at the lower end populated by rarer species, SIVAE overall achieves higher scores. Such behaviour is expected in a more flexible representation learned through our self-supervised pretraining. This flexibility can be seen as beneficial, with yielding softer classification boundaries across taxa can provide more informative measures of classifier uncertainty. BirdNET generalises poorly for these species, over-confidently mis-specifying the inverse of the ground truth, suggesting species are absent when present or present when absent. Some of this behaviour could be attributable to label noise, though generally this indicative of challenges faced by discrimiative models outlined in 1. Scores broken down by species across datasets (see Figure. 11) show most species achieve *>* 0.85 with a small number performing very poorly, likely due to insufficient target (labels) or source (audio) signal or failure of representations to effectively generalise for rare species calls [33]. Top-1 scores are markedly higher than BirdNet for SIVAE, suggesting latent representations are a strong foundation for inter-species discrimination. Both VAE models show some improvement over BirdNET for crowded soundscapes in the UK for certain species (see ‘occlusions only’ in Figure. 2). BirdNET achieves higher AP scores for species that we have very few examples for, likely due to a larger pool of training labels and data augmentation strategies.

To elucidate the comparison in terms of scale: BirdNET’s original model (2021) [2] is a deep residual convolutional network trained for a bespoke set of species on 80,000 labelled examples, with a minimum and maximum of 3500 and 350 samples per label respectively, each corresponding to 3s of audio. Each new species addition requires training a brand new model for all species. We train independent *linear* classifiers on our soundscape prior where each label corresponds to 60s of audio using only a fraction of the training examples. For SO UK, 1620 total examples with 740 to 10 labels, for SO EC, 2161 examples with 280 to 10 labels, and for RFCX 3781 examples with 42 to 26 labels. Ghani et al [4] equally use linear models to fine tune BirdNET and Perch representations, however this an approach requires strong labels to effectively generalise which is not a cost-effective solution for PAM.

BirdNET use many data augmentation strategies to diversity their training set from 80,000 to 1.5 million spectrograms, including temporal shifts, temporal and spectral stretching and audio mixup (randomly sampling and summing waveforms) [2]. In our pipeline we use the following strategies: during VAE inference before fitting classifiers, a single shift by half a frame is applied to each spectrogram to minimise bordering effects from the framing operation, doubling the number of inputs; samples are drawn from the latent distribution during training to introduce some diversity into the training set for classifier training. Our method would equally benefit from similar augmentation strategies as BirdNET, suggesting further potentially significant gains can still be made.

Our proposed approach of having a strong learned prior on soundscapes dramatically improves the efficiency of downstream models, which can be trained in a fraction of the time on a fraction of labelled samples, while providing on-par performance with very large and computationally expensive bespoke models for a specific task - in this case, species detection.

### 2.4 Interrogating shift-invariant latent representations

In this subsection, we evaluate the consistency of learned representations for each VAE variant with respect to the species detection task.

Figure 3 presents the pairwise L2 distances between the most contributory frame embeddings used for prediction when a species was actually present in the audio. Classifier weights are used for feature selection to identify the predictive features and attention scores are used to perform frame selection from 60s audio files. To ensure comparability across models, L2 distances are normalised by the dataset average to account for inter-model latent space diversity. The results show that for most all species, introducing intra-frame invariance to representations results in smaller distances between predictive frames. SIVAE representations are more tightly bound in the latent space, suggesting that during learning the model was better able to focus on spectro-temporal morphology and yield stronger clusters for call types and species.

**Figure 3.**
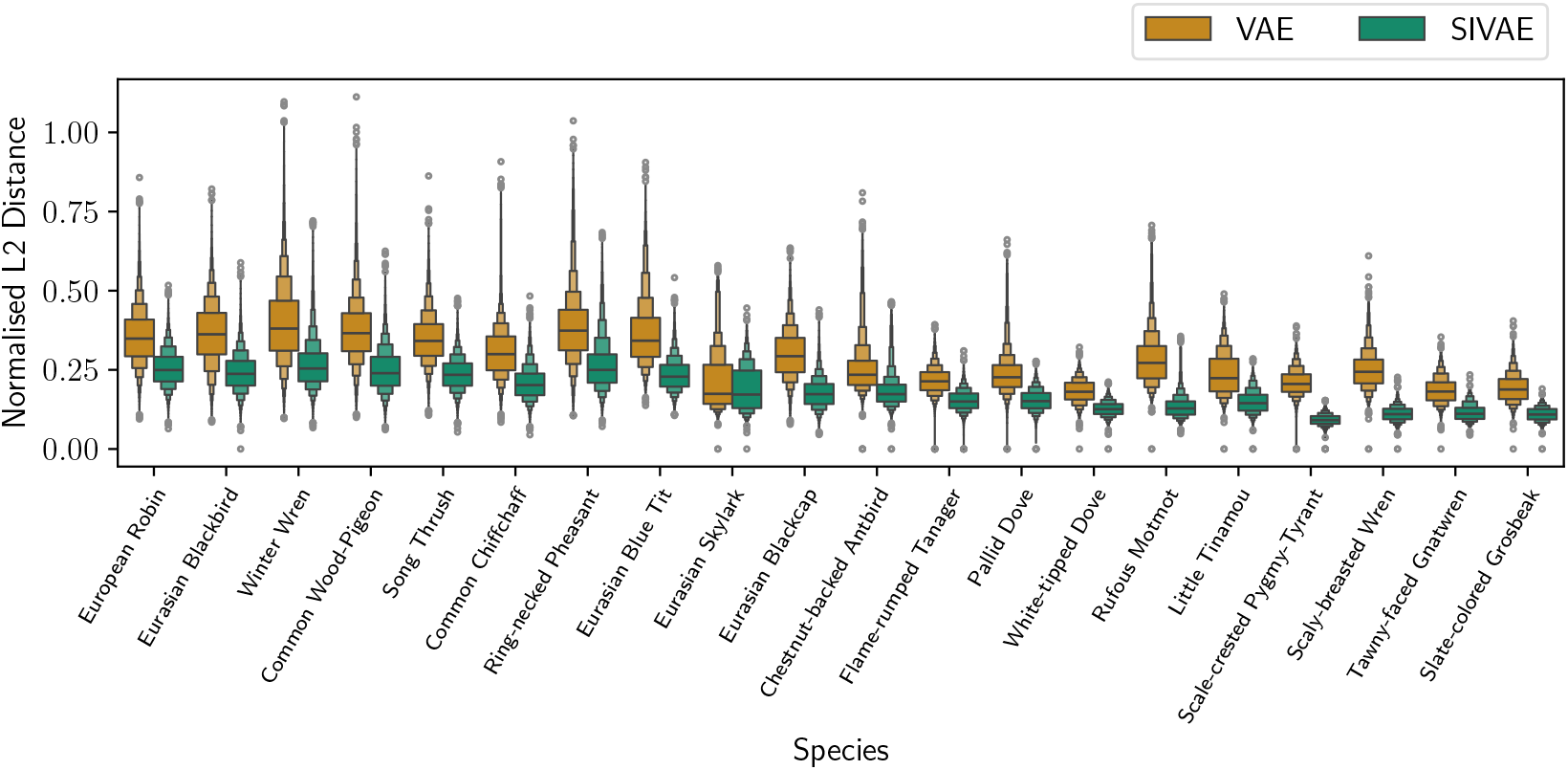
Latent space consistency of frame embeddings between the classic VAE and SIVAE for 10 species from each country in the SO dataset. Pairwise euclidean (L2) distances between the most predictive frame representations where the species was present in the clip shown as distributions. Distances are normalised by the dataset mean for comparability.

For certain avian species in RFCX, distances between representations are much smaller (see Fig. A.2)), for example the Reg Legged Thrush, Elfin Woods Warbler, and Puerto Rican Woodpecker. By cross-referencing their overall scores with the generated species representations shown in Fig. 4, we observe that the classifier has failed to focus on the desired species. This indicates that empty frames were selected before calculating distances, resulting in feature representations much closer to the standard normal prior, effectively describing a silent background. We can see some outliers as such in both SO UK and SO EC where distances are very small. In those cases this indicates a similar failure of selection.

**Figure 4.**
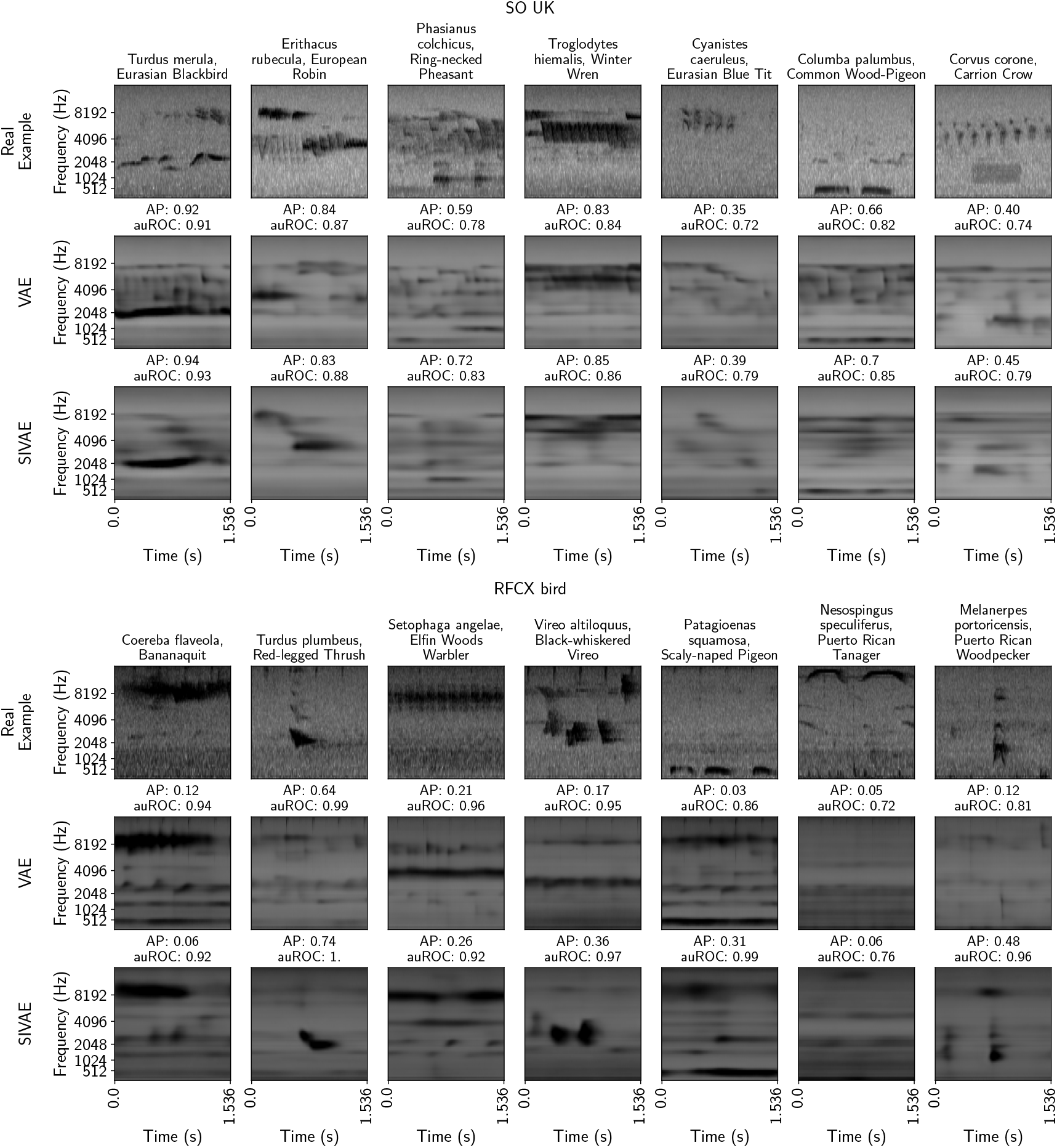
Comparing accuracy and interpretability of proposed model innovations in the SO UK and RFCX avian species. Exemplar spectrogram frames containing calls (1st row) for different species andwgenerated spectrograms (see section Sec. 4.4.3) for two different VAE training paradigms including a classic VAE (2nd row) and a VAE with intra-frame shift-invariance (SIVAE) (3rd row). VAE spectrograms visualise examples of particular latent representations that are the basis of a positive detection, generated by a traversal of the latent space of the generative model as directed by the classifier.

Overall, evidence supports the intuition that an arbitrary temporal co-ordinate system assigned during training has a negative effect on representation consistency. The introduction of shift-invariance improves consistency within species while supporting improved discriminative ability.

### 2.5 Evaluating the impact of novel representation learning algorithm on interpretability

In this section we evaluate the ability to inspect the basis of classification by leveraging the power of our VAE decoder to generate novel samples and reconstruct them as spectrograms. The visualised generative representations are indicative of the key features of a particular species, as learned based on traversal of the latent space directed by a linear classifier. We contrast both the effectiveness and reliability of this method across the VAE and SIVAE model variants.

The introduction of shift-invariance leads to more ecologically relevant specifies classifiers while increasing detection accuracy compared to the classic VAE. Figure 4 demonstrates qualitative differences in the learned generative species representation according to individual specifies classifiers (using the method described in 4.4.3) set against typical call examples. For each dataset real examples (1st row) were selected by identifying common call patterns through manual observation and selecting the most isolated call examples present in the data. Generated species representations can be regarded as spectro-temporal representations of the basis a positive classification. For all reconstructions the classic VAE exhibits finer temporal resolution in the reconstructions while the shift-invariant model has much smoother reconstructions with some spectral banding. This is particularly noticable for the Blue Tit (*Cyanistes caeruleus*) and Winter Wren (*Troglodytes hiemalis*) in 4, where the form of the call is visible but obscured by some spectral banding across the biophonic spectral region (2-6 kHz) due to interpolating across all dimensions relevant for the prediction. The SIVAE tends to be less subject to using spurious correlated soundscape components, resulting in an overall gain in classification performance.

Examining generated species representations from the RFCX bird dataset, shift-invariant samples make significant gains, both in terms of discriminative performance and model interpretability, when interrogating the basis of species detection. For example, the full spectro-temporal morphology of the Thrush (*Turdus plumbeus*), Woodpecker (*Melanerpes portiocensis*) and Vireo (*Vireo altiloquus*) calls become clearly distinguishable in the generated sample. This comes alongside a step change in the average precision of the model. The fidelity and reliability of the learned generative species representation appears related directly to the precision of the classifier. More precise detectors do appear more consistent with respect to the ecologically relevant target information, typically detailing the morphology of specific calls more effectively. However results suggest this does not necessarily mean they are less subject to leveraging spurious correlated soundscape components, as in the case of the Eurasian Blackbird (*Turdus merula*) in SO UK. A well-balanced dataset containing isolated calls in addition to those present and obfuscated by other species calls would likely lead to robust models. Further disentanglement of soundscape components leveraging contextual prior knowledge to factor bird calls into independent arrangement of feature would fundamentally assist in separating source soundscape components. The Puerto Rican Tanager (*Nesospingus speculiferus*) is a significant failure case consistent across both the classic VAE and the SIVAE where the model failed during training to identify relevant calls. Lower auROC and AP during training indicate the classifier failed to effectively fit to latent features on this occasion, through this was not consistent across random initialisations where one model trained on the classic VAE achieved success during the 9 training runs. Further interrogation of network initialisation, hyper-parameter tuning and expanding the training data should yield consistent results. In this case the classification model was unable select the relevant features from latent representations to focus on the spectro-temporal morphology of the relevant call.

Figure. 5 demonstrates a critical failure case when detecting the Puerto Rican Spindalis (*Spindalis portoricensis*). The example calls (first two rows) provided in the left most spectrograms show the initial source for the presence label. Generated species representation shares the same spectro-temporal morphology as the example call in each of these spectrograms, as well as those in the frames leveraged by the classifier (which differ from the original source), providing corroborating evidence that this particular structure is the basis of the detection. However, the final example spectrogram contains a *different call type* emitted by the same species. Most species exhibit a diversity of call types, deploying different vocalisations in different contexts. One of the benefits of self-supervised generative pre-training is this natural diversity should be preserved in the latent space. However some research suggests that generative models such as VAEs may struggle to learn underlying rare generative factors [33]. This particular failure case demonstrates either a failure of the VAE or the classifier due to a skewed distribution of intra-class call variability in the data. The VAE may have failed generalise to more significant variations in vocalisations beyond small variations in the spectro-temporal morphology of this species’ call, resulting in lost signal during the encoding process. Alternatively this may simply be a failing of the classifier. Despite being well described in the latent space, this call is less common and its representation is a significant outlier in in the detection task. The classifier overfits to the more commonly observed features in the training set, selecting inappropriate features for the outlier during inference. We suggest that the more likely scenario is (2) given the labels are infrequent and very weak.

**Figure 5.**
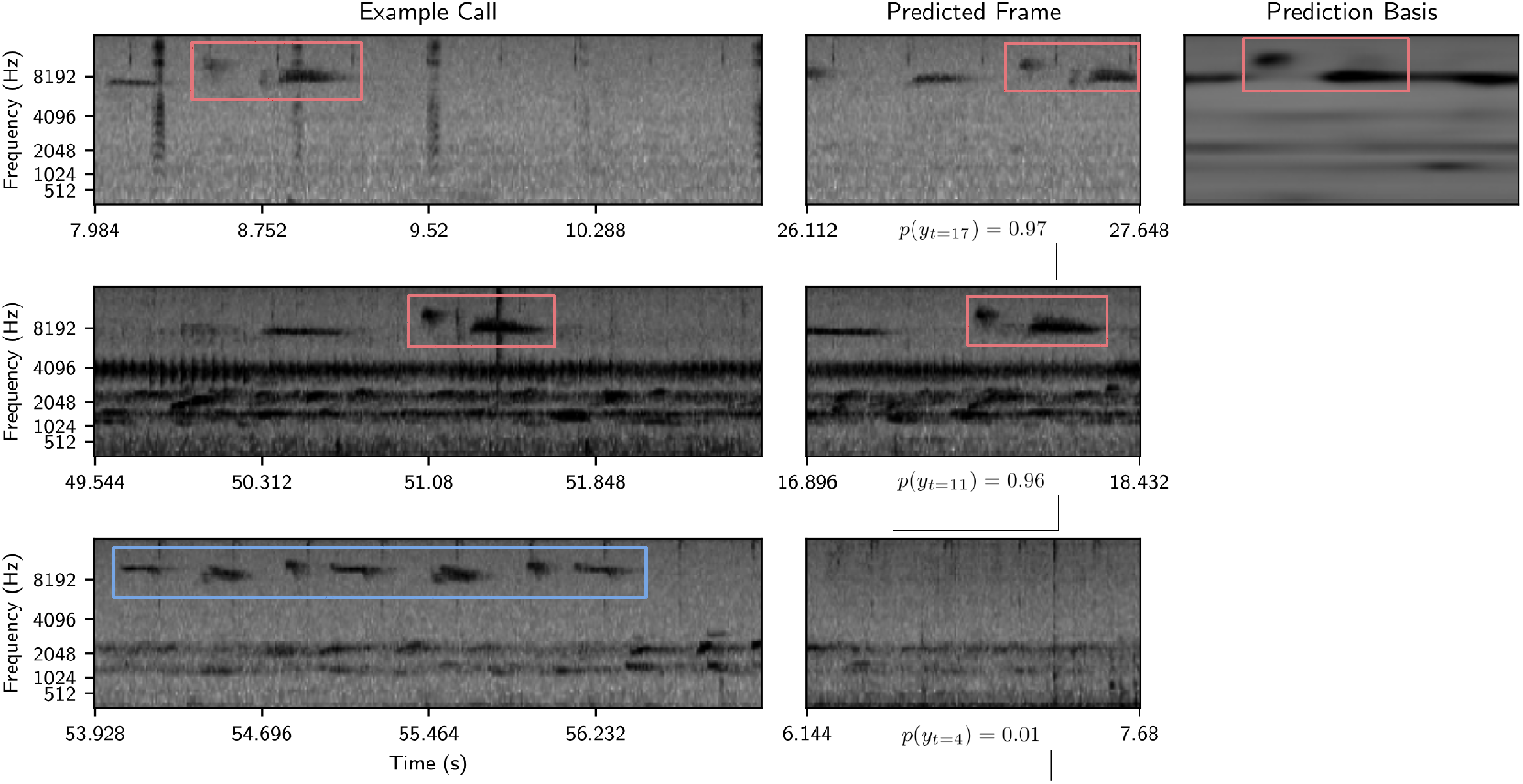
SIVAE detection failure case for the Puerto Rican Spindalis (*Spindalis portoricensis*). **Left**: The spectrotemporal morphology of calls of a consistent type are shown in rows 1 and 2 (highlighted in red), while a different type is shown in row 3 (highlighted in blue). **Centre**: The first call type is correctly identified with high probability (rows 1 and 2) while the second call type results in a very low probability (row 3). **Right**: Decoded generated species representation describing the basis for the detection is consistent with the first call’s spectro-temporal morphology. A red box is drawn around the comparable shape to emphasise the similar call component.

## 3 Conclusion

In this study, we introduced a novel self-supervised deep learning pipeline for modelling complex soundscapes as a prior for a bioacoustic detection task to help create cost-effective generalisable and interpretable models with a high efficiency of human effort. Specifically, our proposed approach used disentangled Variational Auto-encoders to learn a strong prior on whole soundscapes for use in a downstream species detection task in tropical and temperate terrestrial ecozones. Due to observed limitations in the descriptive ability for such a task as a result of the auto-encoder algorithm embedding an arbitrary temporal co-ordinate system into learned representation, we introduced a novel innovation for the soundscapes, intra-frame shift invariance, to help improve consistency, accuracy and interpretability of representations used in the detection task. Quantitative experiments show our approach was both effective in terms of discriminative performance, achieving the highest area under the ROC (0.92) and Top-1 accuracy (0.67), outperforming BirdNET V2.4 (0.89, 0.52) while achieving good average precision despite using very weak presence annotations for each minute of audio.

More importantly, our qualitative experiments demonstrate a new method for interpreting the basis of a positive species detection using generative species representations learned by individual species classifiers. Experimental results highlight the effectiveness and interpretability of VAE latent representations for avian species detection. By performing latent space traversals of the VAE latent space directed by a linear classifier, generated spectrograms demonstrate the reliability of species detection models, highlighting spectro-temporal structures used for making predictions and emphasising examples where spurious correlated soundscape components are used in the detection process. We performed a comparative analysis between a classic VAE and a intra-frame shift invariant VAE, exploring success and failure cases and conducting an analysis of the structure of the latent space, demonstrating this class of models yields more compact representations and results in improved species detection scores.

Nevertheless, every approach has its limitations. Firstly, while introducing shift-invariance aids consistency of the representation and improves discriminative performance, this sacrifices some of the reconstruction fidelity we achieve in the classic VAE. Some of this could be alleviated by optimising the training strategy and ensuring the shift-invariant model observes more truly unique examples as well as shifted versions. However enforcing structure onto a compression necessarily results in some degradation in representational capacity. Secondary generative models could be developed to help refine reconstructions while preserving the representational power developed in this work. Secondly, due to the circularity of the translational prior, under certain initial conditions shift invariant VAE models did not effectively ascertain the task. Further work could examine expanding the cross-decoding scheme to ensure the model is forced to explain those examples using the shift prediction network. Thirdly, despite VAE representations effectively describing call diversity, classifiers can still fail on a weakly labelled prediction in significant cases where species with high diversity of call types are not observed frequently enough, resulting in over-fitting. Finally, one limitation of using a learned generative representation from a binary presence absence task is we cannot easily realise a diversity of species representations. The species presence transformation can be applied at different co-ordinates in the latent space, but knowing where to start is one of the challenges.

Future work should establish ecological generalisability, tune for long tail distributions and enhance computational efficiency. Preliminary experiments using a class-balanced loss [34] and class-weighted sampling during training yielded mixed results, though there is significant space for further exploration, and both methods are supported in our code. There are many more opportunities to enhance the interpretability of a VAE for soundscape analysis by embedding task-specific prior knowledge to structure the latent space. Further experiments examining methods to structure features by leveraging the temporal dynamics of biophonic events may help disentangle representations providing additional gains when predicting species presence while providing sematic interpretability. To meet the challenges of the biodiversity crisis, the transformative potential of machine learning offers a powerful tool to meet the needs of rapid data collection of PAM. It is paramount we build data efficient tools that are both accurate and critically interpretable to support robust ecological inference and advance the science as a whole.

## 4 Methods

This section begins by briefly outlining our audio processing. The details our adjustment to the VAE architecture are introduced, outlining the implementation of intra-frame shift invariance. Following this, the classification model for species detection and the training methodology are outlined in detail. Finally the interpretability mechanism for inspecting the basis of a positive species detection is described.

### 4.1 Pre-processing

To enable comparison with BirdNET each one minute audio file is transformed following [2] to yield a log-Mel spectrogram **x** (full details of FFT and Mel transformation parameters are also available in our prior work [28]). Frames within our model at all scales have a temporal resolution of 1.536 seconds. Full contiguous mel spectrograms are a maxmimum duration of 59.904 seconds for equal framing.

### 4.2 Designing intra-frame shift-invariant representations

Learned audio representations typically describe a short fixed window (frame) of a spectrogram, which can be aggregated by averaging [35], histogramming [28] or directly used as features for a prediction task, such as species detection. These types of features enable temporally specific representations of the audio, but the process of framing introduces an arbitrary coordinate system regarding the absolute position of an acoustic event within a frame, see Fig. 1A and 1B for an illustration of a single frame under several translational shift 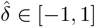 with circular boundary conditions. As each of these frames contains the same content, we would like a learned representation to be disentangled with respect to the temporal coordinate system of the frame.

Fig. 6 illustrates our proposed model for encoding latent representations from a spectrogram. Shape information in the spectro-temporal domain is modelled locally at different scales using a deep residual convolutional neural network [36] ℰ_CNN_ to an intermediate feature map, which is temporally framed. These framed features are processed through two distinct multi-layered perceptrons: 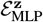 which produce an intra-frame shift-invariant latent representation (**z**) of that frame of audio; and 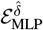, which predicts the intra-frame shift 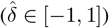 of the frame contents, with respect to a learned canonical timeline. These models are learned through a generative process, based on an adapted variational auto-encoder, and draws on related work in computer vision [30] disentangling texture and pose.

**Figure 6.**
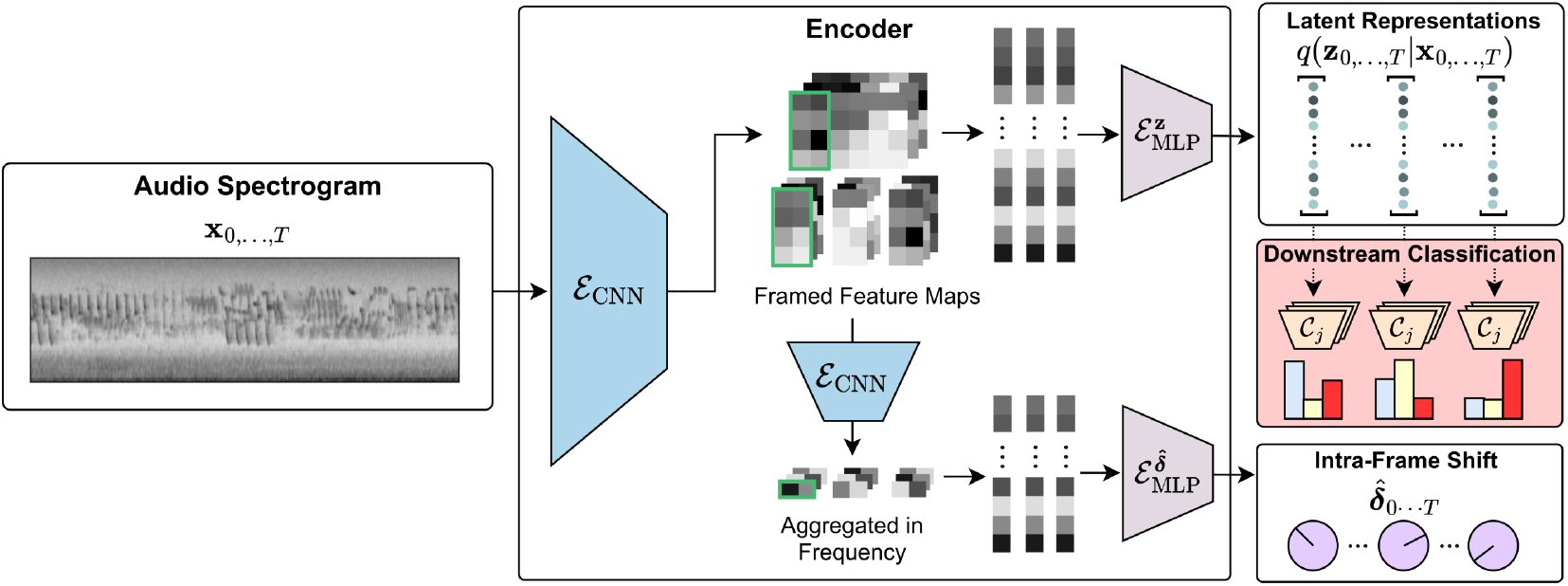
Encoding architecture overview. Spectrograms **x**_0,…,*T*_ are encoded into intra-frame shift-invariant latent representations, **z**_0,…,*T*_, with a corresponding shift 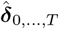 for each frame. These representations are learned in a fully self-supervised manner, see section 4.2 for details; subsequently, classifiers can be trained using **z** as a feature.

#### 4.2.1 Learning shift-invariant latent representations

We formulate the task of disentangling a learned representation of absolute temporal position as a self-supervised learning problem. We sample transformed instances of each observation **x**^′^ by applying a translation 𝒯(**x**, *δ*) drawn from an appropriate prior distribution *δ* ~ *p*(*δ*). We train a *shift prediction* neural network 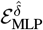 (Fig. 6) conditioned on convolutional feature maps to infer a maximum-a-posteriori estimate for *p*(*δ*|**x**) ∝ *p*(**x**|*δ*)*p*(*δ*). The *content* network 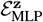 conditioned on the same information infers a full approximation to the posterior over soundscape content *q*_*ϕ*_(**z**|**x**) by variational inference. We use 2-way cross-decoding across the set of instances {**x**, 𝒯(**x**_0_, *δ*_0_), …, 𝒯(**x**_*T*_, *δ*_*T*_)} to ensure translation information is not included in the soundscape content descriptor **z** and temporal variability is funnelled into the alignment network. We train a decoder network, 𝒟, to maximise the likelihood of each instance given the average of the framewise latent representations, 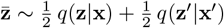 and the predicted alignment factors, i.e. 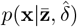 and 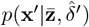. During decoding the inferred alignment factor is used to translate the partially decoded feature representation, via linear-interpolation, before being fully decoded back into spectrogram space. During training, at each iteration the model observes spectrogram sample sets consisting of a full contiguous original **x**_0,…,*T*_ and a spectrogram consisting of concatenated independently translated frames of the same spectrogram 𝒯(**x**_0_, *δ*_0_), …, 𝒯(**x**_*T*_, *δ*_*T*_), where *T* denotes the length of the sequence of frames, as illustrated in Fig. 6. Translations are sampled independently from a zero-centered Gaussian prior *δ* ∼ 𝒩 (0, *σ*_𝒯_) and bounded on the interval [−1.0, 1.0] using the modulus operator.

To compute alignment factors 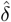, additional down-sampling is computed across the frequency dimension using a (1 ×*f*) convolution and passed through a bottleneck 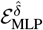 to output a single 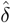 for every frame. To compute the latent content, **z**, feature maps are passed through a second bottleneck 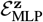 to output an 128-dimensional Gaussian distribution, with diagonal covariance, for each frame. During decoding, we sample from the average Gaussian distribution 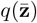 using the reparameterisation trick. 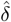 is applied as a translation mid-way through the decoder as in [37]. Note that due to the periodic boundary condition, frames have piece-wise discontinuity at the boundary, which can result in minor inconsistencies when re-combining independently aligned segments into a contiguous sequence of spectrogram frames. To reconstruct a contiguous original **x**_0,…,*T*_, framed feature maps are recombined at the penultimate convolutional layer, while representations used to reconstruct independently translated frames remain discontiguous.

#### 4.2.2 Loss

We use a Gaussian likelihood on the log-Mel spectrogram values given the predicted spectrogram from the decoder.

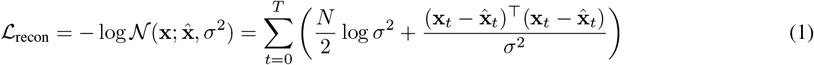

Where 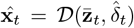 *N* is the number of pixels in a frame and *t* indexes the *T* frames. *σ*^2^ is a shared variance across the dataset, which balances the importance of reconstruction fidelity against the smoothness of the latent representation, where smoother latent representations tend to be more useful in downstream tasks [27]. Practically, we experimented with *σ*^2^ ∈ {0.2, 0.3} as well as inferring a global value and found 0.3 delivered sufficiently good reconstructions to discern spectrotemporal features while still leading to meaningful decodings when interpolating between latent representations.

As the alignment factor 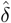 is circular, inference is ill-posed. We regularise learning to predict 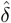 by specifying a zerocentered prior distribution:

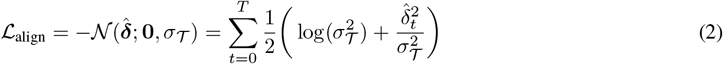

The total loss for our proposed model is the sum of each components as defined in Eq 1 and Eq 2, plus the standard VAE regularisation term, the KL divergence between the approximate posterior with a standard normal prior:

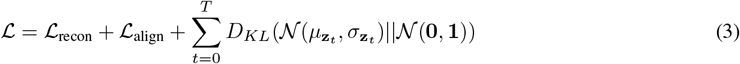

### 4.3 VAE training details

We consider 2 VAE variants: Classic, which is trained with the ELBO loss [26] and Shift Invariant, which optimises Eq. 3. We train 3 separate VAEs of each variant with initial seeds for each dataset. VAE models were trained for 180,000 batches on batches of 6 log-Mel spectrograms of duration 59.904s with 64 mel bins. We optimised the learning rate experimentally by observing validation performance *η* = 0.00004 and use a cosine annealing schedule with linear warmups restarting every 60,000 batches. At training time the model (as detailed in Fig. 6 and Fig. 7) is provided audio-clips of duration 59.904s, resulting in 39 x 1.536s frames that make up the latent time-series.

**Figure 7.**
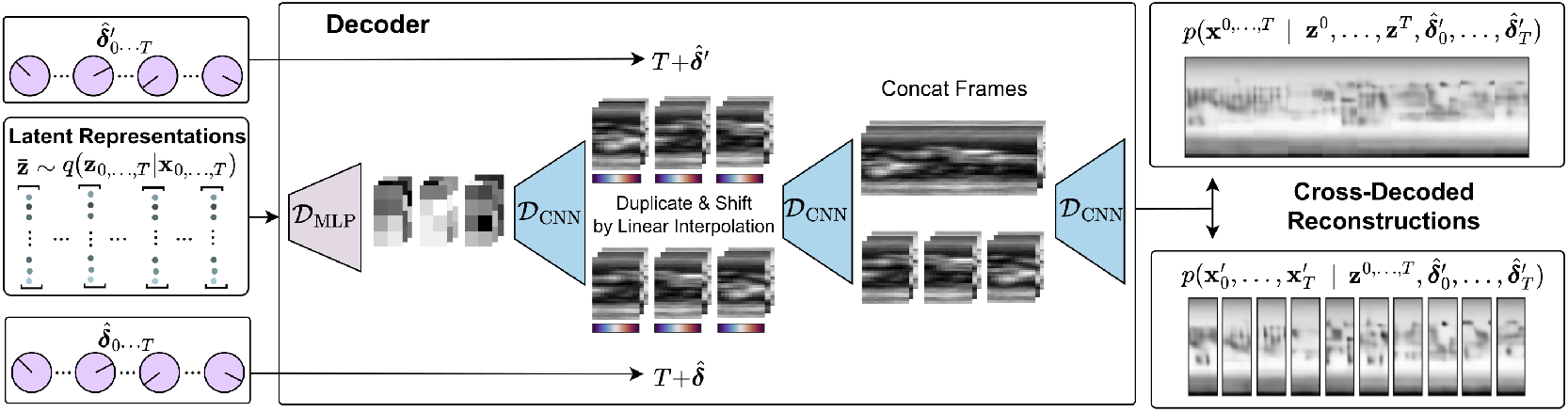
Decoding architecture overview. Samples are drawn from the average Gaussian distribution using the parametrisation trick. The decoder maps the sample **z**^*t*^ to approximate each input spectrogram, both a full contiguous clip and independently translated frames. During decoding each frame the decoded representation is duplicated for each target and is independently shifted in time by applying a translation 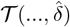

Theoretically the optimal prior for the circular boundary condition is a uniform distribution. However we found experimentally that a normal prior initialised with a small variance and annealed over the first 40,000 batches to approach a uniform distribution reduced the negative impact of attempting to align observations with significant noise or very little foreground signal early in the training process. We smoothly transition *σ*_𝒯_ across the interval [0.01, 2.0] using a bounded sigmoid function. Specifically, let *i*^curr^ denote the current batch index, *i*^min^, *i*^max^ denote the start and ending batch index for the annealing process, 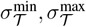 are the minimum and maximum values for the standard deviation of the prior distribution, we set the current value 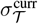 at each batch:

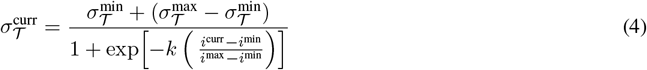

where *k* is the slope of the sigmoid which we set to 1.

### 4.4 Detecting species using weak labels

Acoustic events generated by a species can occur multiple times within each 60s audio sample. However weak annotations denote only a sample-level binary category describing species presence or absence. Drawing on the work of Ilse et al. [38], we formulate the problem of identifying the key frames that trigger a presence prediction as a supervised multiple instance learning (MIL).

#### 4.4.1 Frame-wise logistic regression

We discard the temporal continuity of embedded latent features and instead regard each sample **z**_0,…,*T*_ as a bag of independent instances. In this setup, predictions are made at the instance level and a final bag-level label is computed by aggregating instance level predictions using a permutation-invariant pooling operator. For each species, we train an L1 regularised linear classifier, 𝒞, to predict the probability of a species vocalisation in each frame. Output probabilities are then pooled using an aggregation function pool_*t* ∈ [1,…*T*]_ to extract a bag-level probability. As in [38], we experimented with 3 pooling methods to aggregate frame-level probabilities for the bag-level prediction: (1) the max operator; (2) the mean operator; and (3) a weighted sum, whereby each frame’s weight is learned as a function of itself using a small feed-forward neural network for each species:

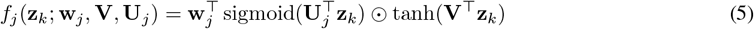

where **z**_*k*_ is a bag instance, **w**_*j*_ ∈ ℝ^*hJ*^, **U**_*j*_ ∈ ℝ^*J*×*h*×*d*^ **V** ∈ ℝ^*h*×*d*^ are learned parameters and ⊙is the element-wise multiplication operator. Each latent representation is further reduced down to a *h*-dimensional feature space using **V**. The gating function **U**_*j*_ learns a species-specific feature selection of the *h*-dimensional embedded signal, while **w**_*j*_ maps the gated signal down to a single value for each instance *t*. A weight for each frame and species is computed by normalising outputs across the bag using the softmax activation function. Finally, these are multiplied with the classifier output probabilities and summed to obtain a final bag-level presence prediction (6).

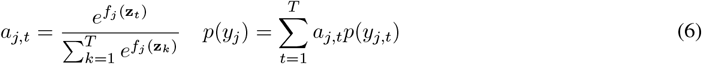

We minimise the binary cross-entropy loss with L1 weight regularisation to sparsify the weights of the classifier for feature selection. For attention models we regularise the weights of the *f*_*j*_ to prevent overfitting using an L2 penalty.

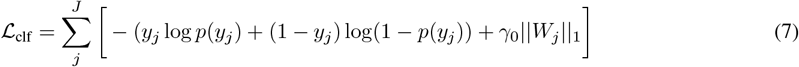

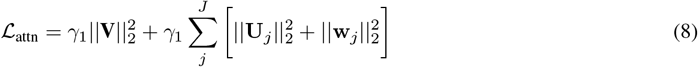

The final loss for the proposed classification model is the sum of each components as defined in Eq 7 and Eq 6:

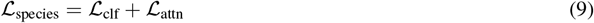

#### 4.4.2 Training and hyper-parameter optimisation

Classifiers for each species were trained concurrently as a unified model. For the SO dataset, there is no overlap between species for each country, therefore the training data was split by country to prevent confusion by fitting to species that will never be present. The species list for each dataset was restricted to the intersection of labels between the training and test set. The training data consists of 60s spectrograms embedded twice by the VAE: the first instance remains unchanged, the second is shifted by half a frame (0.768s) to minimise the effect of signal loss due to the arbitrary boundaries of the frame operation applied during the encoding process. To expandwthe variability in the dataset and inject some noise to improve model robustness, we draw samples from the VAE’s approximate posterior distribution during training. During evaluation we draw *N* = 100 samples and average the output probability to minimise the effect of sample noise. The final input for each classifier is a bag of 78 x 1.536s frames, each with 128 features. For the feed-forward gated attention network, we set the hidden layer *h* to 10. The attention network has a fixed shared 1280 parameters (**V**) with additional 1290 parameters (**U**_*j*_, **w**_*j*_) for each species added, while each classifier consists of 128 parameters.

We performed a 5-fold cross-validation across the 3 pre-trained VAEs of each model class (initialised with different random seeds) and selected the optimal combination of 4 hyper-parameters for each dataset: L1 penalty (*γ*_0_), L2 penalty (*γ*_1_), classifier learning rate (*η*_0_ and attention network learning rate *η*_1_). Performance of each classifier is evaluated using the area under the receiver operating characteristic curves (auROC) and the average precision (AP) for each species. Model selection proceeded by taking the maximum of the sum of the average AP and auROC scores across species and random seeds for each pooling method. We found that optimal pooling method were either max or a learned weighted sum. However reliable and consistent hyper-parameter selection was challenging using max. Output probabilities at the beginning of training and thus the learning trajectory are particularly subject to small differences in weight initialisation, resulting in high inter-model variation in evaluation metrics. For the attention mechanism the weights are initialised to a uniform distribution [39] and the model is able to smoothly learn to adjust the weights by backpropagation to identify relevant frames.

To evaluate detection performance, we adopted three commonly used score metrics: (1) the average precision (AP) evaluates model precision at every threshold by calculating the area under the precision-recall curve; (2) the area under the receiver operating characteristic curve (auROC) evaluates the ratio of TPR to FPR at every threshold, providing a summary measure of model sensitivity; (3) Top-1, or recall at K, is a measure of accuracy in a multi-class prediction task, calculated by ranking probabilities and evaluating the proportion of correct detections.

#### 4.4.3 Interpreting learned representations using generative models

We take advantage of the generative potential of the VAE decoding mechanism [40] to inspect the basis of classification, providing a means to interpret learned representations. We generate examples of spectrograms by decoding samples from regions of the latent space where a species is classified as present. As introduced in [28], we first define a presence/absence transformation for each species *s* in the latent space using each linear classifier. Specifically, we apply an affine transformation 𝒮_*j*_(**z**_*k*_) to traverse the hyperplane of each species-specific classifier:

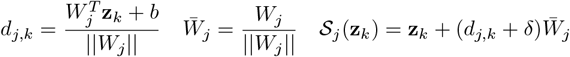

where **z**_*k*_ is a sample from an appropriate starting region of the latent space, *d*_*j*_, *k* is the signed distance to the hyperplane for species *j* and 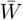 is the direction normal to the hyperplane. *δ* is a tunable distance parameter beyond the decision boundary controlling the amplitude of the signal in the generated spectrogram. This transformation performs a linear interpolation of values in the VAE latent space, changing a classification from absent to present. With the VAE decoder we generate reconstructions to identify spectro-temporal signals that correspond with each transformation 𝒮_*j*_(**z**_*k*_). This provides a means to visually inspect a decoding of the representations that the classifier used to predict each species. While this can technically be applied at any point in the latent space, a good starting point is an approximation to a generic average embedding. For SO we take the average of latent features within each site, and determine which site to use for a given species according to where it was most commonly observed. For RFCX we take the average for each taxonomic group (birds / frogs), excluding embeddings that are known to contain the other taxonomic group. Reconstructions of average embeddings for each site are shown in Fig. 8.

**Figure 8.**
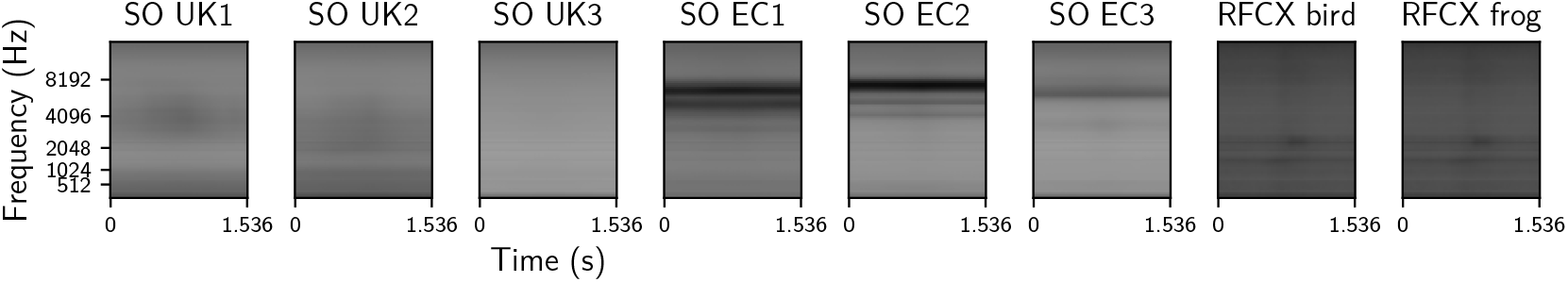
Generated spectrograms of samples classified as absent for all species. These are used as the start point for applying species presence transformations 𝒮_*j*_(**z**_*k*_) to generate species calls and thereby inspect inspect the basis of the classification. Samples **z**_*k*_ are calculated by taking the average of latent feature embeddings with respect to a specific subset. For SO, each habitat has quite different spectral profiles and species populations, so wedefinethesubset as the habitat average. For RFCX we take the average across each taxonomic group.

## APPENDIX

### A.1 Demonstrating shift invariance

**Figure 9.**
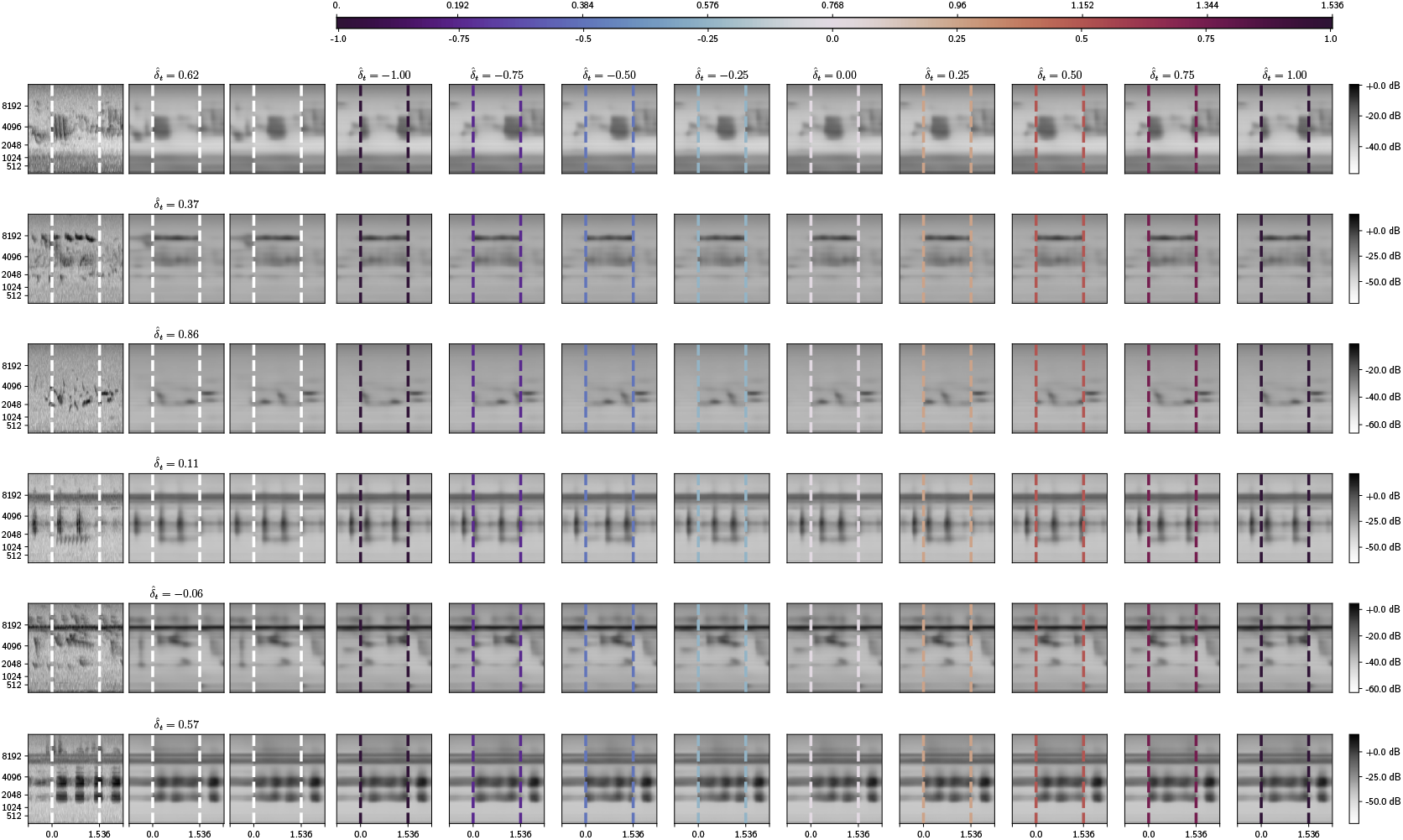
**First column**: A sample from each site in the SO dataset rendered as the original spectrogram. White dashed lines indicate the 1.536s frame being considered. **Second column** Reconstruction 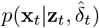 using the alignment factor inferred by the shift prediction network 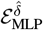 **Third column** A prototype frame reconstruction can be generated by setting the alignment factor to zero. **Remaining columns**: Control over absolute position in time can be parameterised by sampling the alignment factor 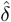 from a continuous and periodic temporal co-ordinate space (top) to instantiate different shifts of the prototype sound event. Dashed line colour corresponds to the value of the alignment factor 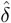. A circular wrap around can be observed when signal falls off the denoted 1.536s period, while signal beyond these boundaries remains unchanged.

### A.2 Distance between Species Embeddings

The following plots show pairwise distances between frames the with highest probability of a species after applying attention weights for examples where the call was present. These are broken down by dataset to show the overall difference in representation consistency for each VAE model class.

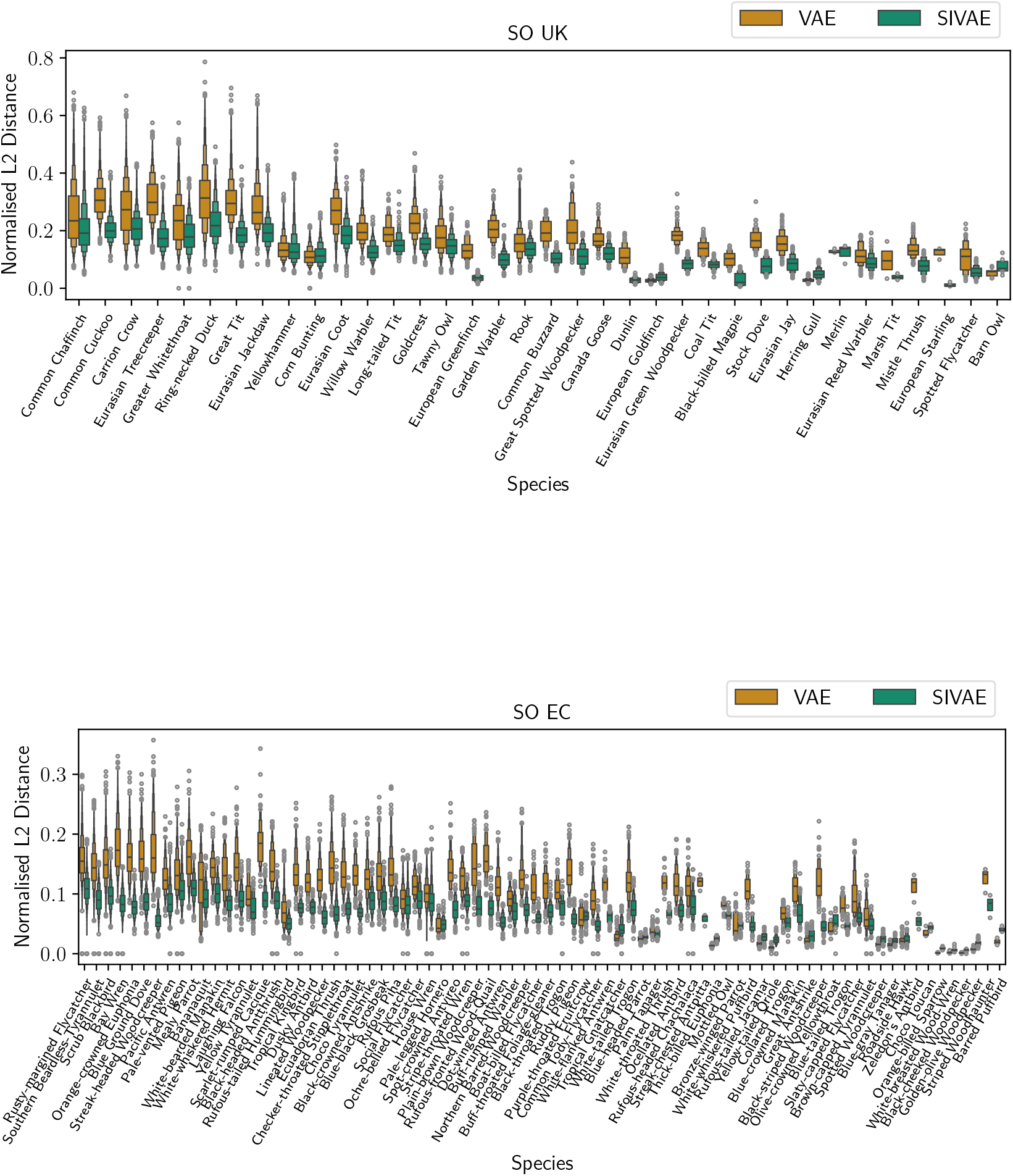
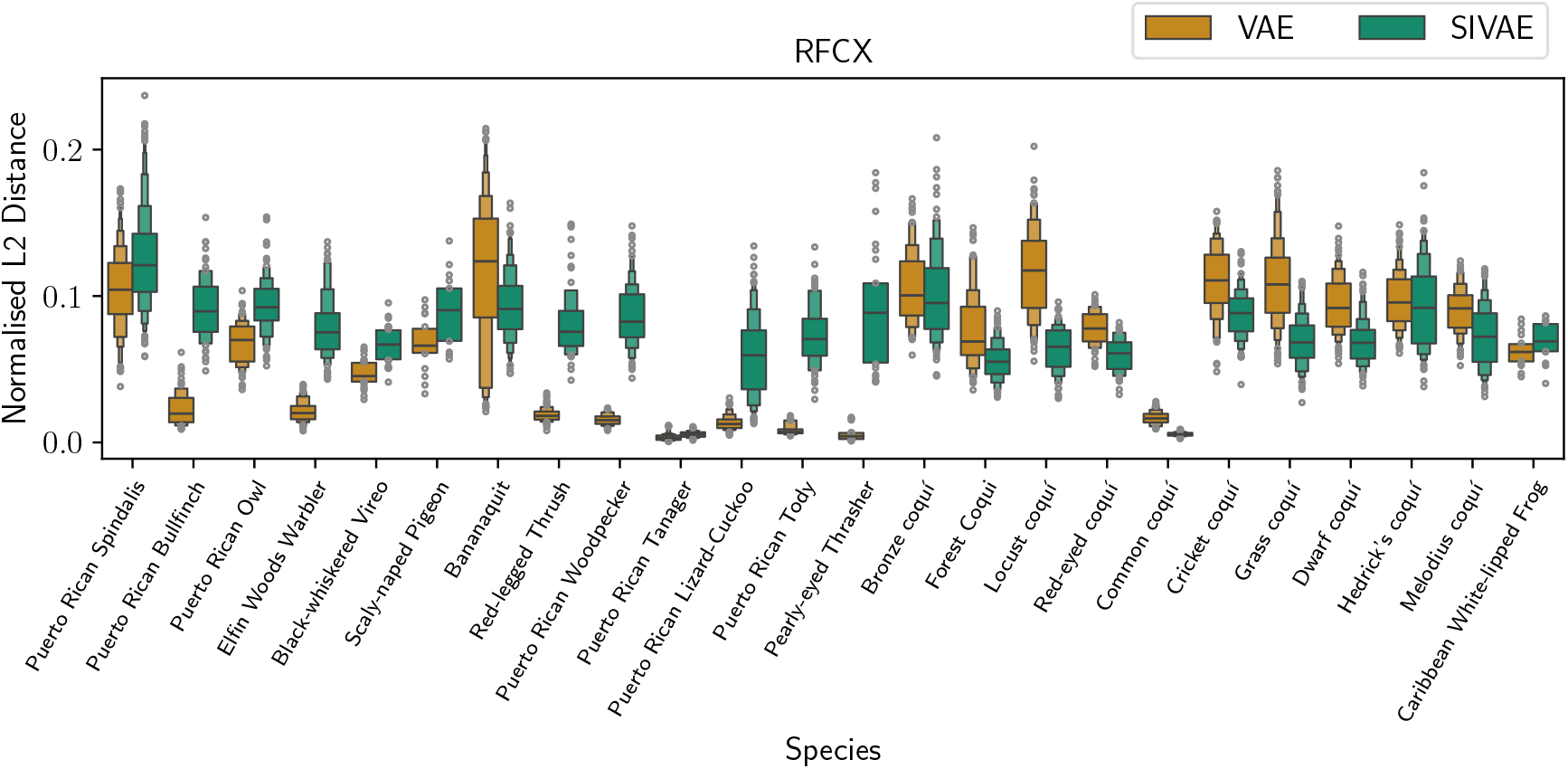

### A.3 Relationship between label count and average precision

Model average precision as a function of the count of presence training labels for each species, broken down by dataset. Achieving a high positive predictive score on species detection tasks is species dependent at the tail end and will be determined by the distinctiveness of the call and the uncertainty in the VAEs latent description.

**Figure 10.**
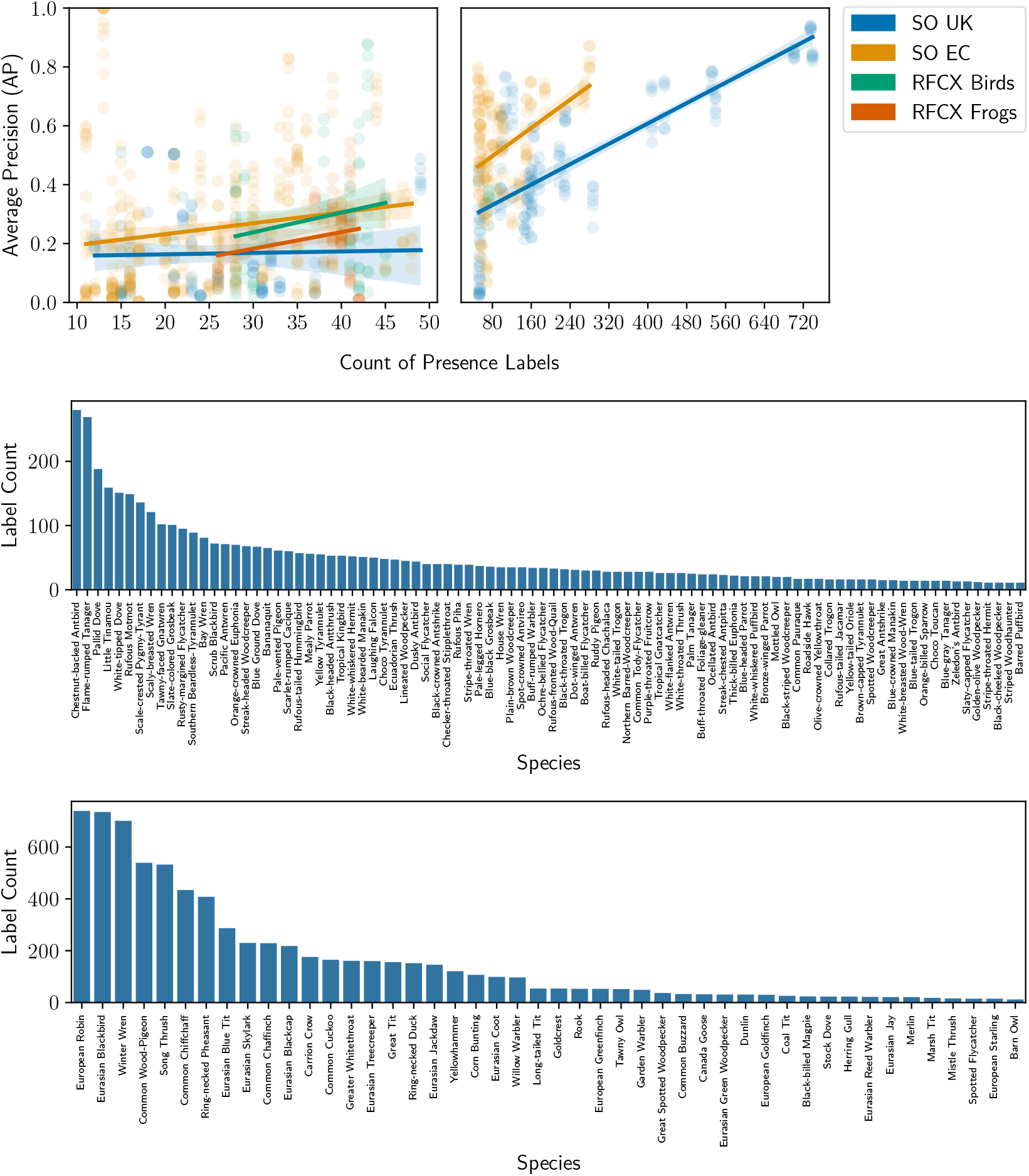
Fewer than 50 labels for each species typically results in low precision, though for many species we are still able to achieve good scores. Beyond 50 labels the trend shows a linear increase, noting that there are fewer species with this many labels. Label counts for each species in the SO dataset.

### A.4 Breakdown of scores by species and dataset

Box plots show the average precision (AP) and area under the receiver operating characteristic (ROC) curve (auROC) across species for all models.

**Figure 11.**
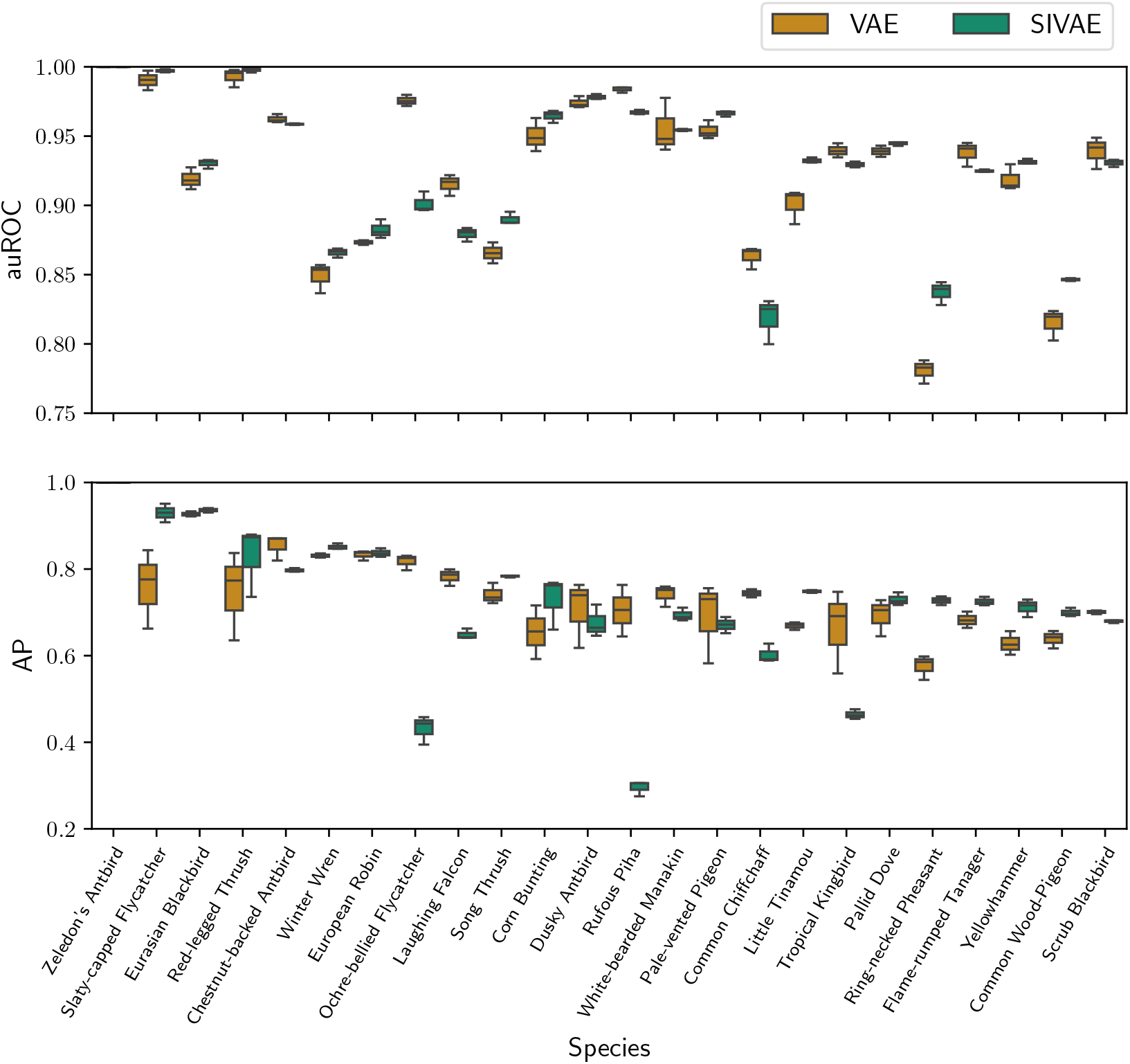
A breakdown of the distribution of scores for the top 24 species across datasets for VAE and SIVAE, sorted by training label frequency in descending order. Further breakdowns for each dataset are available in the appendix A.4.

**Figure 12.**
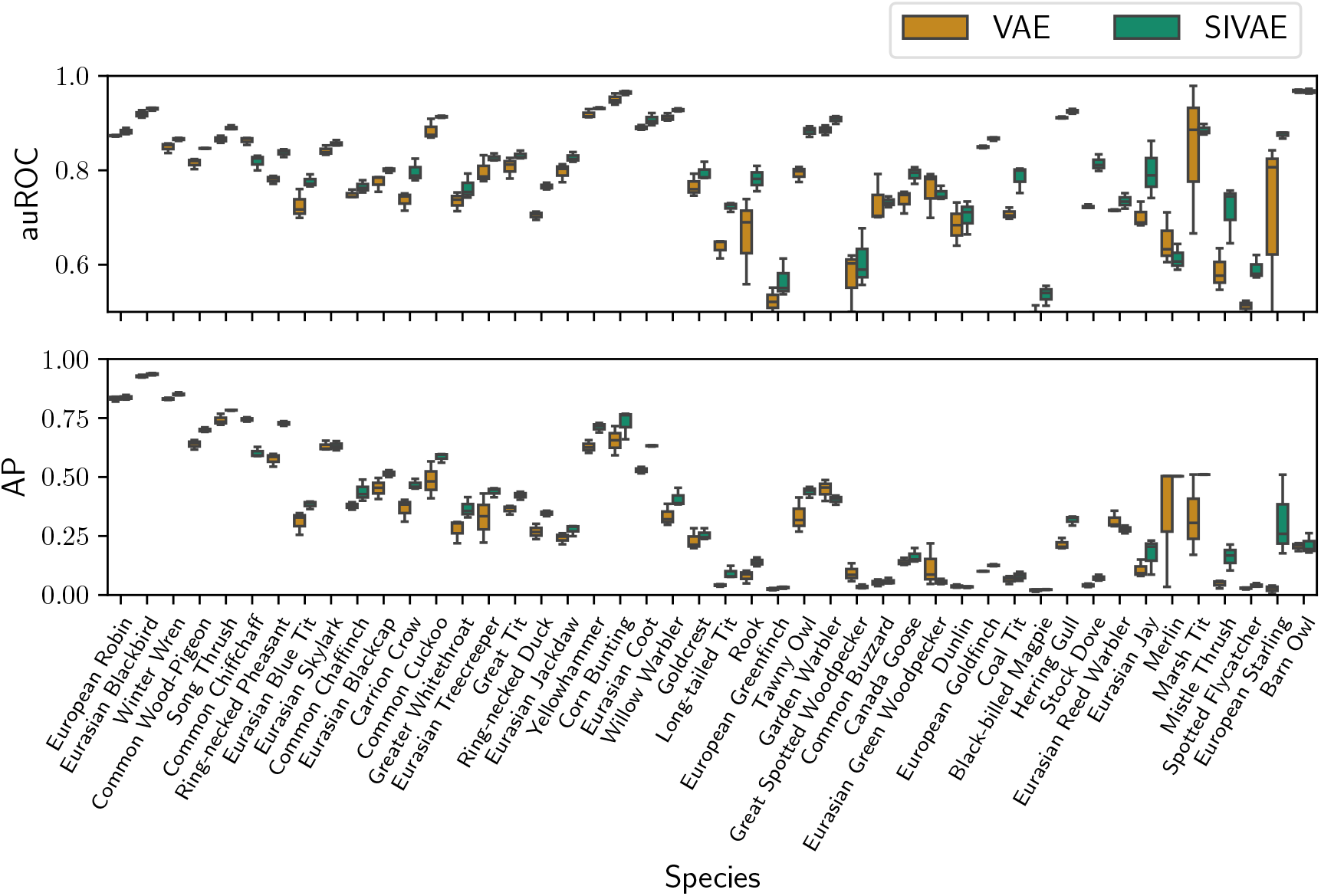
Per-species distributions of the AP and auROC for the SO UK dataset.

**Figure 13.**
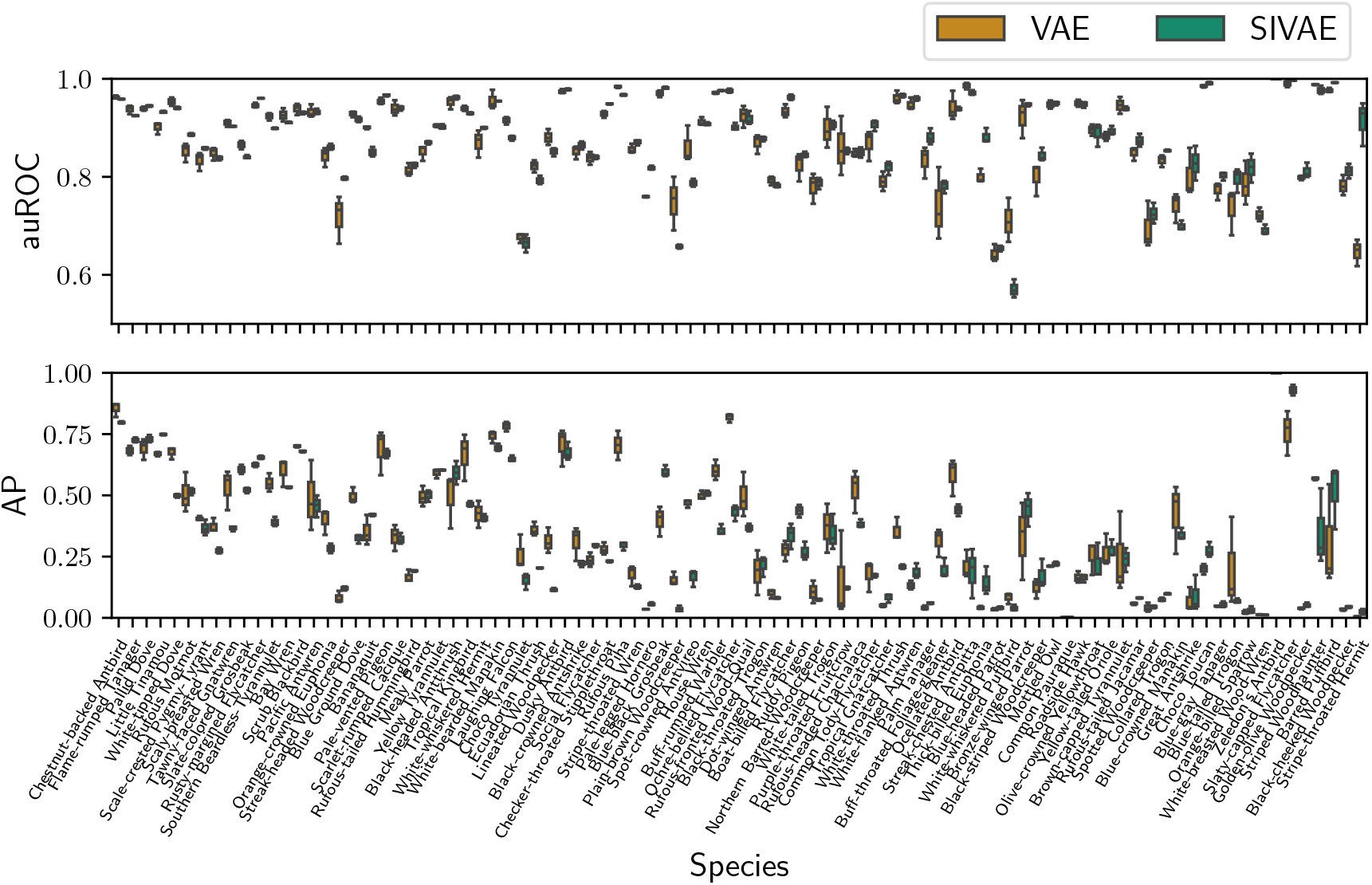
Per-species distributions of the AP and auROC for the SO EC dataset.

**Figure 14.**
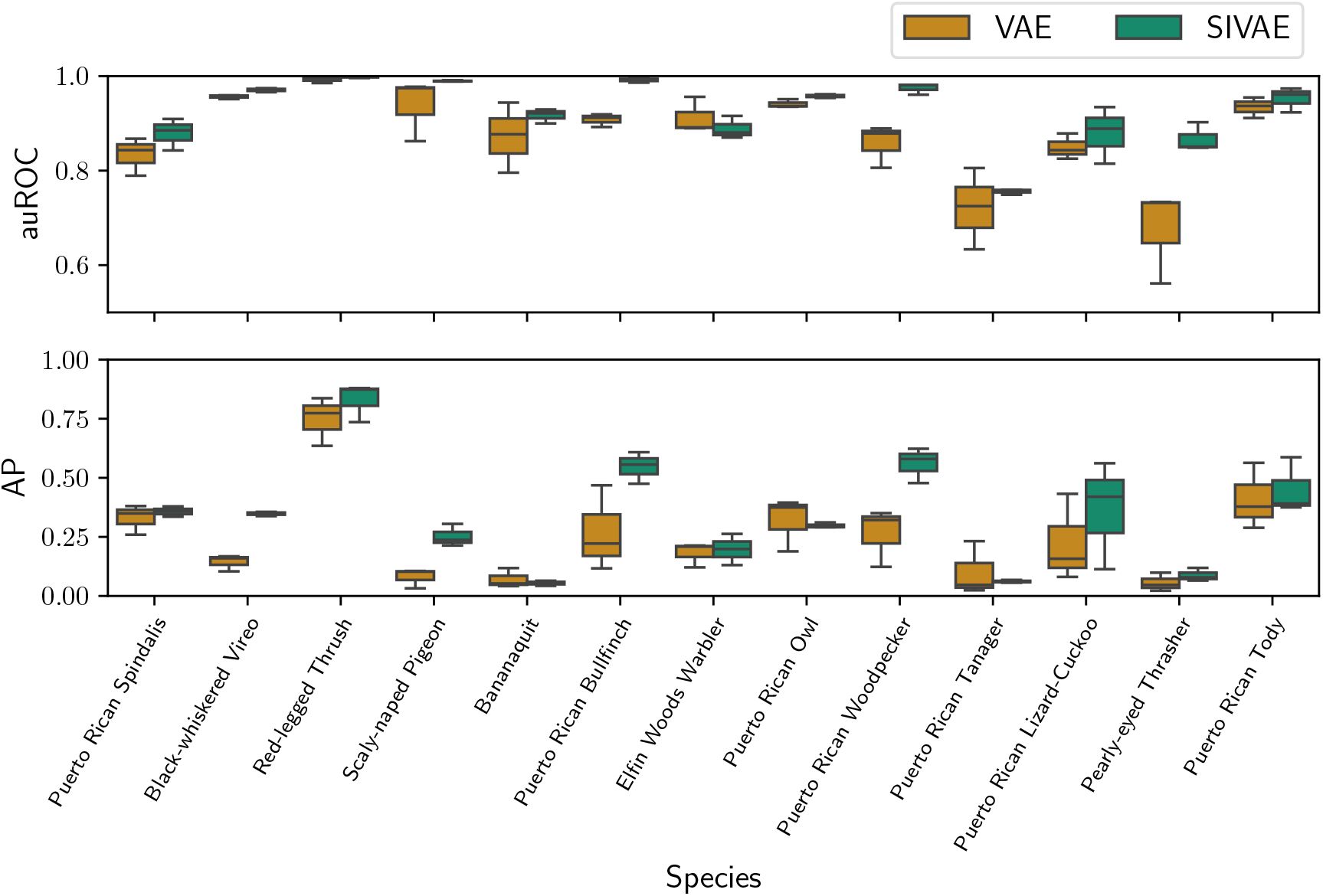
Per-species distributions of the AP and auROC for the RFCX bird dataset.

**Figure 15.**
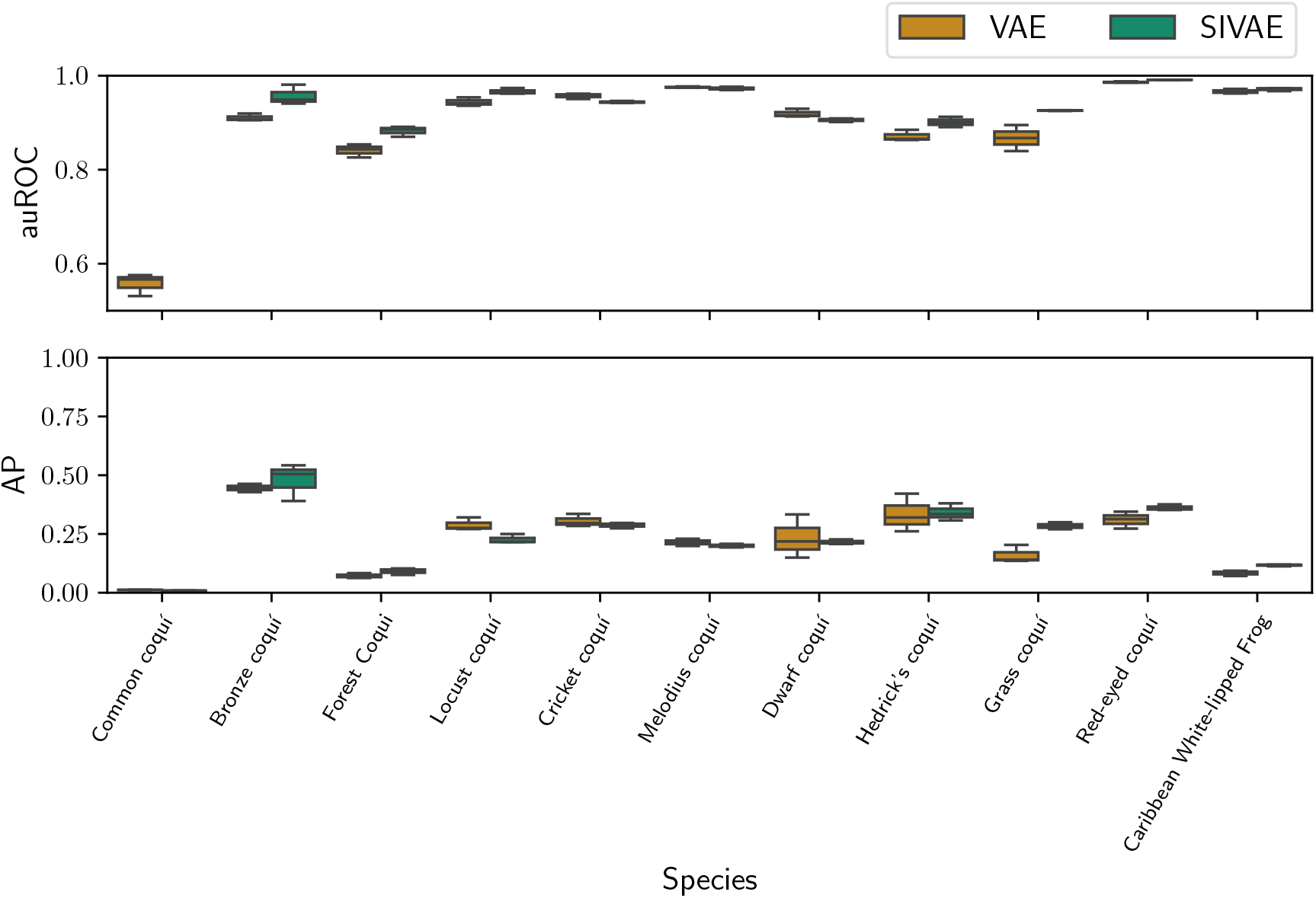
Per-species distributions of the AP and auROC for the RFCX frog dataset.

### A.5 Generated Species Representations

Additional generated species for the other examples the SO and RFCX datasets, including Ecuadorian species, and frogs from Puerto Rico.

**Figure 16.**
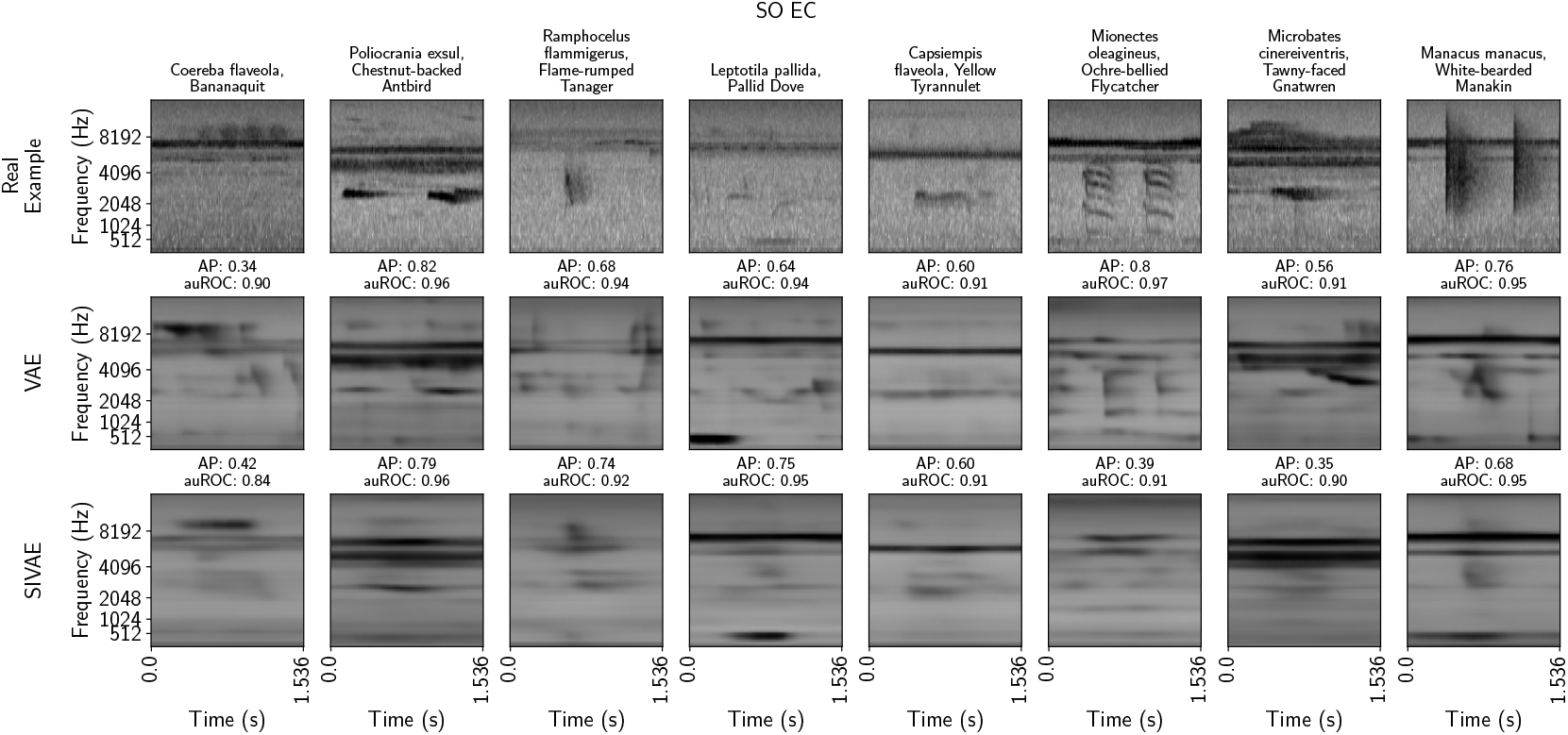
Comparing real (top row) and generated examples (bottom two rows) by the two VAE variants for the SO EC dataset.

**Figure 17.**
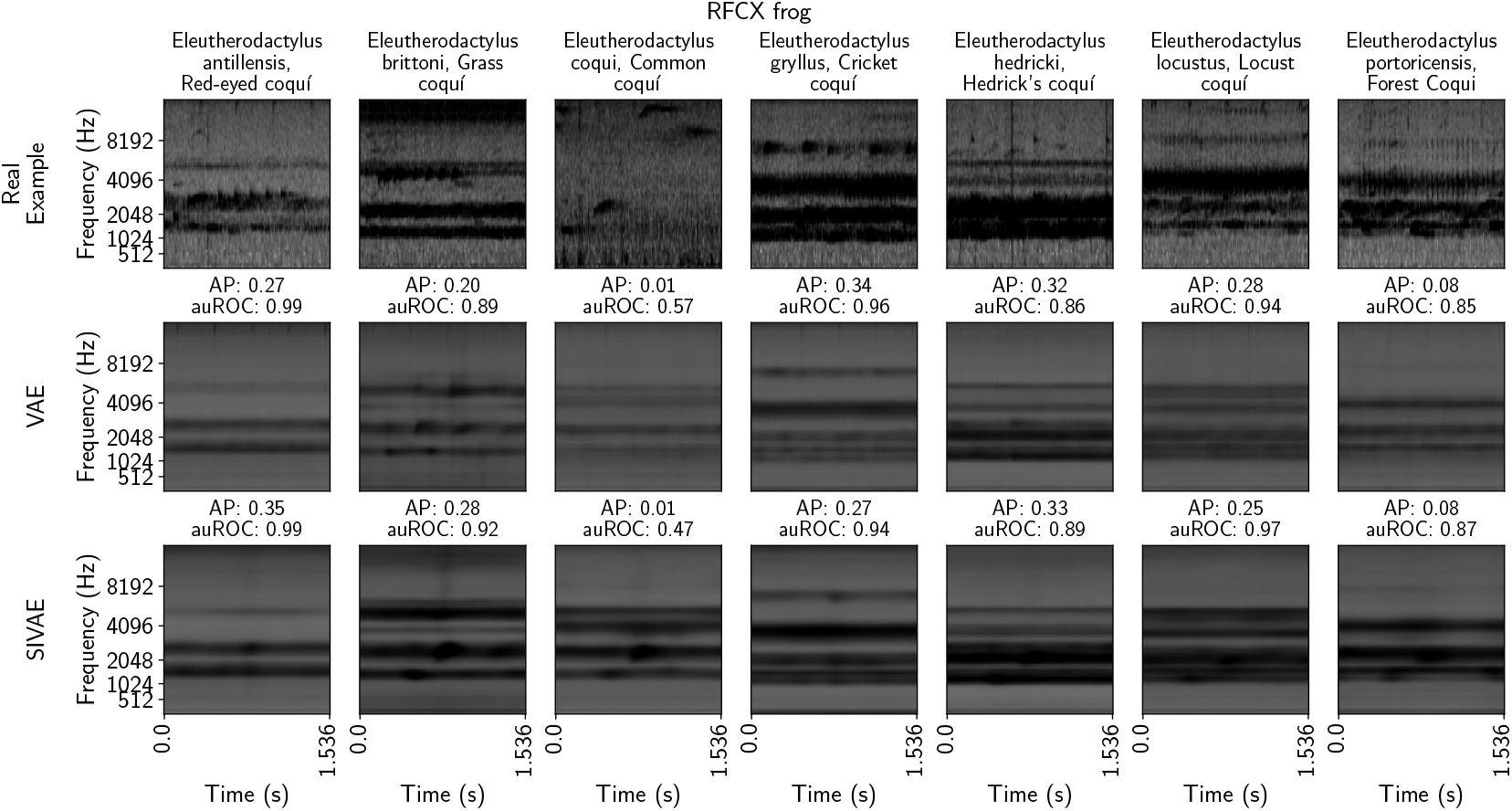
Comparing real (top row) and generated examples (bottom two rows) by the two VAE variants for the RFCX frog dataset.

